# Piccolino regulates the architecture of the ribbon at cochlear inner hair cell synapses

**DOI:** 10.1101/2022.12.15.520589

**Authors:** Susann Michanski, Rohan Kapoor, Anna M. Steyer, Wiebke Möbius, Iris Früholz, Frauke Ackermann, Mehmet Gültas, Craig C. Garner, F. Kent Hamra, Jakob Neef, Nicola Strenzke, Tobias Moser, Carolin Wichmann

**Author notes:** Equal contribution (Susann Michanski, Rohan Kapoor). Shared correspondence (Tobias Moser, Carolin Wichmann).

## Abstract

Cochlear inner hair cells (IHCs) form specialized ribbon synapses with spiral ganglion neurons that tireless-ly transmit sound information at high rates over long time periods with extreme temporal precision. This functional specialization is essential for precise sound encoding and is attributed to a distinct molecular machinery with unique players or splice variants compared to conventional neuronal synapses. Among these is the active zone (AZ) scaffold protein piccolo/aczonin, which is represented by its short splice variant piccolino at cochlear and retinal ribbon synapses. While the function of piccolo at synapses of the central nervous system has been intensively investigated, the role of piccolino at IHC synapses remains unclear. In this study, we characterized the structure and function of IHC-synapses in piccolo gene-trap mutant rats (*Pclo^gt/gt^*). We found a mild hearing deficit with elevated thresholds and reduced amplitudes of auditory brainstem responses. Ca^2+^ channel distribution and ribbon morphology were altered in apical IHCs, while their presynaptic function seemed unchanged. We conclude that piccolino contributes to the AZ organization in IHCs and is essential for normal synaptic transmission.

## Introduction

Ribbon synapses are involved in vertebrate vision, hearing and balance. They are specialized in terms of function, morphology and molecular composition to enable indefatigable neurotransmission over long time periods. Depending on the ribbon synapse type, the electron-dense appearing synaptic ribbon can tether up to several hundreds of synaptic vesicles (SVs) (Matthews and Fuchs, 2010; Moser et al., 2019; Wichmann and Moser, 2015). In addition to employing the ribbon-specific protein RIBEYE, that constitutes the main component of the synaptic ribbon (Becker et al., 2018; Jean et al., 2018; Maxeiner et al., 2016; Schmitz et al., 2000), the molecular composition of ribbon-type AZs shows further differences from conventional neuronal synapses (Chakrabarti and Wichmann, 2019; Moser et al., 2019). For example, ribbon synapses employ different isoforms of presynaptic proteins such as rab3-interacting molecule 2 (RIM2) instead of RIM1 (Grabner et al., 2015; Jung et al., 2015a). Moreover, the presynaptic multi-domain protein piccolo/aczonin, is represented solely by its short splice variant piccolino as shown for photoreceptor (Regus-Leidig et al., 2013) and cochlear inner hair cell (IHC) ribbon synapses (Butola et al., 2017; Michanski et al., 2019; Regus-Leidig et al., 2013).

The function of piccolo at conventional synapses has been investigated intensely (Cases-Langhoff et al., 1996; Fenster et al., 2003; Gundelfinger et al., 2015; Leal-Ortiz et al., 2008; Mukherjee et al., 2010). Piccolo together with bassoon is involved in synapse assembly, SV clustering and maintaining synapse integrity (Gundelfinger et al., 2015). Piccolo seems especially important for the SV pool organization. At the calyx of Held, piccolo was found to organize the readily-releasable vesicle pool (RRP) (Parthier et al., 2018) and at the endbulb of Held, piccolo is required for normal SV replenishment (Butola et al., 2017). Furthermore, a reduction of SVs, specifically from the total recycling pool was found at hippocampal neurons lacking piccolo (Ackermann et al., 2019). In contrast, piccolino’s role at ribbon synapses remains poorly understood. Importantly, in comparison to full-length piccolo, piccolino lacks a number of interaction sites for presynaptic binding partners such as CAST/Munc13 and RIM, L-type Ca^2+^ channels as well as the interaction site for its homologue bassoon (Regus-Leidig et al., 2013). Instead, piccolino was found to interact with the ribbon component RIBEYE (Müller et al., 2019), in line with its localization exclusively at the synaptic ribbon (Dick et al., 2001; Limbach et al., 2011; Michanski et al., 2019; Regus-Leidig et al., 2013). While at conventional synapses, the functions of piccolo and bassoon seem partially redundant (Altrock et al., 2003; Gundelfinger et al., 2015; Leal-Ortiz et al., 2008; Mukherjee et al., 2010; Waites et al., 2013), piccolino’s function at ribbon synapses might be unique and could differ from piccolo’s function at conventional synapses. Piccolino’s absence or reduction had a striking structural impact on photoreceptor ribbon synapses: the altered ribbon shape went along with a reduced number of SVs around the ribbons (Müller et al., 2019; Regus-Leidig et al., 2014). A previous study of the auditory system reported that piccolino KO mice lack functional deficits on the level of auditory brain-stem response (ABR) thresholds (Li et al., 2021).

Given the visual phenotype and the prominent piccolino expression in IHCs, we performed a comprehensive structural and functional study on piccolo gene trap mutant rats (*Pclo^gt/gt^*) (Ackermann et al., 2019; Medrano et al., 2020; Müller et al., 2019) by combining systems and cell physiology with confocal, stimulated emission depletion (STED) and electron microscopy. We observed an altered Ca^2+^ channel distribution and 3D reconstructions from electron microscopic data uncovered changes in ribbon morphology for a subset of synapses, resulting in two morphologically distinguishable ribbon categories in *Pclo^gt/gt^* IHCs. Category 1 ribbons appeared completely normal, while category 2 encompassed small, spherical ribbons that lacked SVs at their upper ribbon part. Recording ABRs, we discovered a mild hearing phenotype despite our findings of normal IHC Ca^2+^ currents and exocytosis. Our data suggest that piccolino is involved in the proper formation of synaptic ribbons, likely via organizing RIBEYE, and potentially via the ribbon, normal clustering of Ca^2+^ channels. We conclude that piccolino is essential for normal organization of ribbon type AZs and required for normal hearing.

## Results

### Reduced ABR wave I amplitudes and elevated ABR thresholds in *Pclo^gt/gt^* rats

To investigate the role of piccolino in the auditory system, we commenced our study by recording ABRs in piccolino-deficient (*Pclo^gt/gt^*) rats at the age of two months. We found a significant amplitude reduction of ABR wave I, which represents the compound action potential of spiral ganglion neurons (SGNs) to 73% of normal (1.68 ± 0.08 µV, N_animals_ = 10 for *Pclo^gt/gt^* rats *versus* 2.38 ± 0.16 µV, N_animals_ = 10 for *Pclo^wt/wt^* rats (*P* < 0.01, two-way repeated measures ANOVA with post-hoc Holm-Šidák correction for multiple comparisons) at 80 dB (peak equivalent, pe, 20 Hz clicks) (Fig 1A, B). The successive ABR waves were also slightly reduced in amplitude, but the difference was significant only for wave IV (*P* < 0.05, t-test, Fig 1A). ABR thresholds in response to tone bursts showed a mild but significant elevation for middle and high sound frequencies (approximately 10 - 20 dB; *P_12kHz_*, *P_16kHz_* and *P_32kHz_* < 0.01, two-way repeated measures ANOVA followed by Holm-Šidák multi-comparison test), but not for low frequency tones and broadband click stimuli (Fig 1C). We also measured cochlear amplification mediated by outer hair cells by recording the distortion product otoacoustic emissions (DPOAEs) which were unaltered (Fig 1D). Overall, we report a mild but significant impairment in synchronous sound onset responses of SGNs despite intact cochlear amplification, which is consistent with a sound encoding deficit at the first auditory synapse upon genetic disruption of piccolino. This hearing phenotype appears more pronounced than that seen in piccolo mutant mice with maintained piccolino expression in IHCs (Butola et al., 2017) or that of RIBEYE KO mice (Becker et al., 2018; Jean et al., 2018).

**Figure 1.**
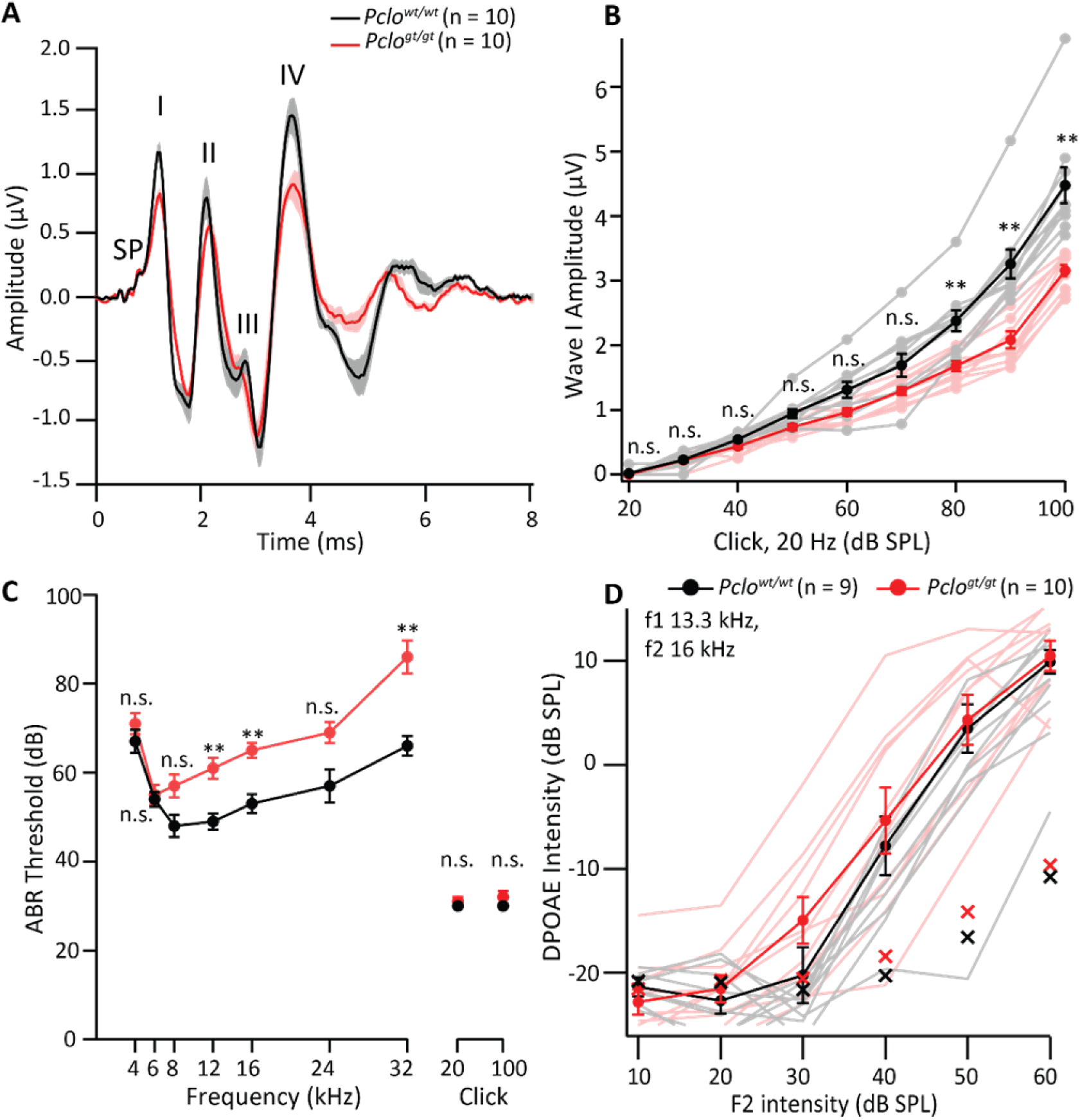
Reduced wave I amplitudes and elevated thresholds of auditory brainstem response. **(A)** Average ABR waveforms in response to 80 dB (pe) clicks (20 Hz) in 2 months old rats (N_animals_ = 10 each for *Pclo^wt/wt^* and *Pclo^gt/gt^*). Lines represent averages, shaded area represents ± SEM. SP: summating potential (hair cell receptor potential), roman numerals (I – IV): ABR waves generated along the auditory pathway. **(B)** ABR wave I amplitudes in response to 20 Hz clicks at different sound pressure levels show a significant decrease in *Pclo^gt/gt^* rats at 80dB and above, implying impaired synchronous sound evoked firing of their SGNs, (*P* < 0.01, two-way repeated measures ANOVA followed by post hoc Holm-Šidák multi-comparison correction). Dark lines represent averages, lighter ones represent individual traces. **(C)** *Pclo^gt/gt^* rats show elevated ABR thresholds for middle and high frequency tone bursts, whereas thresholds in response to low frequency tone bursts and click stimuli (applied at 20 and 100 Hz stimulation rate) appear comparable; *P_12kHz_*, *P_16kHz_*, and *P_32kHz_* < 0.01, two-way repeated measures ANOVA followed by Holm-Šidák multi-comparison correction. **(D)** DPOAE amplitude in response to pairs of simultaneous sine waves (f_1_ = 13.3 kHz, f_2_ = 16 kHz) at increasing sound pressure levels (intensity of f_1_ is 10 dB above f_2_) appear comparable in *Pclo^wt/wt^* (N_animals_ = 9) and *Pclo^gt/gt^* rats (N_animals_ = 10), implying unaltered cochlear amplification. Dark lines represent mean ± SEM, lighter ones represent individual traces, crosses represent noise floor. Asterisks indicate significance levels with \**P*<0.05, \*\**P*<0.01, \*\*\**P*<0.001.

### Smaller ribbons but larger PSD areas at *Pclo^gt/gt^* active zones

We next investigated ribbon synapse abundance and morphology by performing immunohistochemistry on whole-mounted apical portions of the organ of Corti acutely dissected from 2 months old *Pclo^wt/wt^* and *Pclo^gt/gt^* rats, followed by confocal and super resolution STED microscopy. We first used an antibody that recognizes a peptide corresponding to amino acid 2012 to 2351 of rat piccolo, designed to bind to full-length piccolo as well as the shorter splice variant piccolino. In *Pclo^wt/wt^* organs of Corti, we observed immunofluorescent puncta at every afferent synaptic contact, colocalizing with the synaptic ribbon (labeled with an antibody against CtBP2/RIBEYE), as well as at efferent synapses (immunofluorescent spots not colocalized with ribbons). *Pclo^gt/gt^* organs of Corti, on the other hand, showed complete absence of any piccolo/piccolino-specific immunofluorescence (Fig 2A, B). Next, we performed triple-labeling using antibodies against CtBP2 (labeling the synaptic ribbon), bassoon (the presynaptic active zone (AZ)) and homer1 (the postsynaptic density). The localization of the synaptic markers seemed unaltered. We observed that the total number of synaptic ribbons (total count of CtBP2 spots) and the number of anchored ribbons (number of CtBP2 and bassoon juxtapositions) stays intact upon piccolino disruption (Fig 2D). CtBP2/piccolo co-labeled sections as shown in Fig 2B were included for estimation of total ribbon counts.

**Figure 2.**
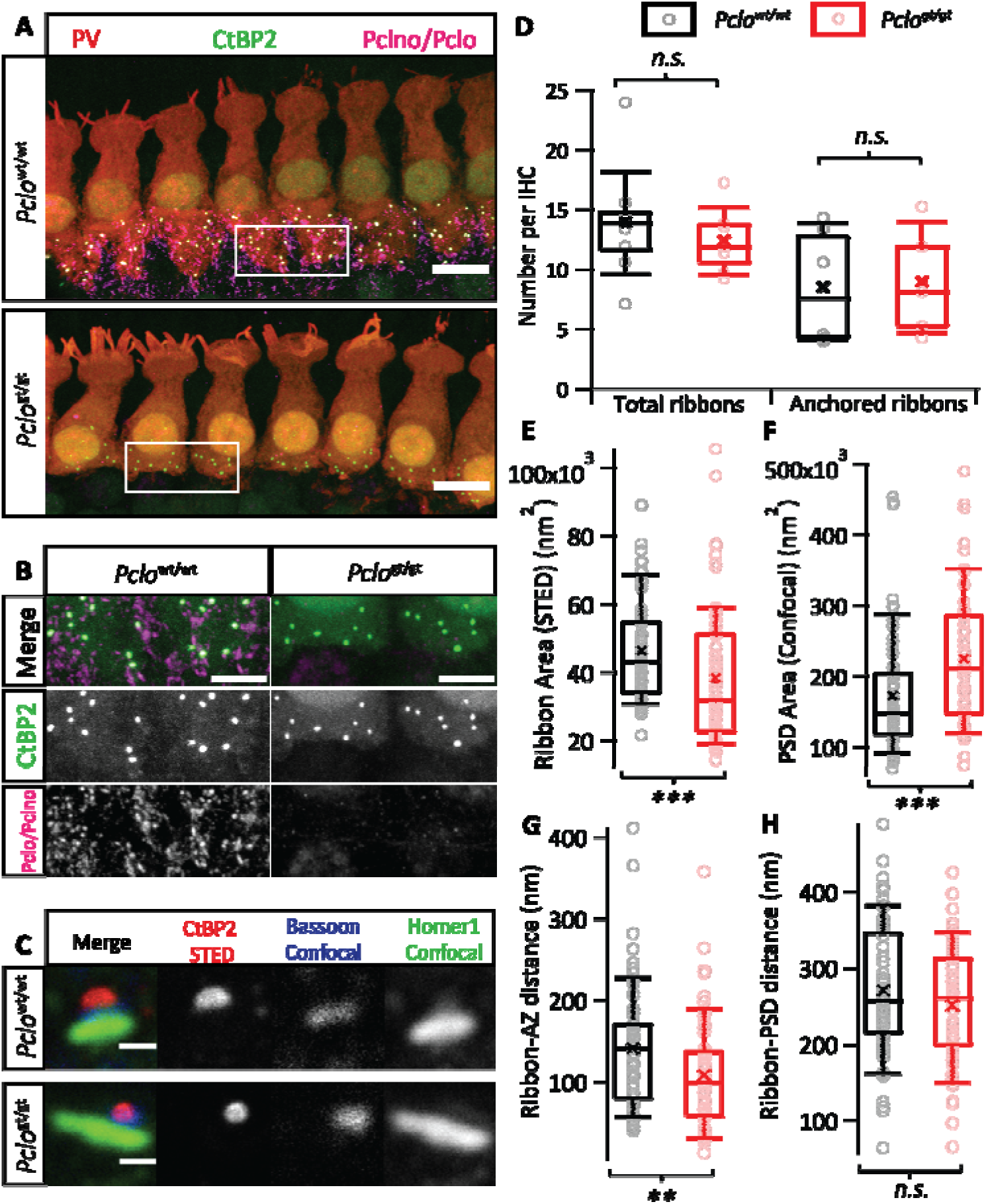
Absence of piccolino results in smaller synaptic ribbons and larger PSDs. **(A)** Maximal projections of confocal sections of apical organs of Corti from 2 months old *Pclo^gt/gt^* and *Pclo^wt/wt^* rats. The whole mounts were co-labeled for IHC cytosol marker parvalbumin (PV, red), synaptic ribbon marker CtBP2 (green) and piccolo/piccolino (magenta). Scale bars = 10 μm. **(B)** Zoom-ins from insets in (A), (here removing the parvalbumin-channel for clarity) highlight the absence of piccolo-specific immunofluorescence puncta in and around *Pclo^gt/gt^* IHCs, in contrast to colocalizing CtBP2-piccolino spots for *Pclo^wt/wt^* IHCs and efferent piccolo immunofluorescence (not colocalising with the synaptic ribbon). Scale bar = 5 μm. **(C)** Triple labeling of homer1 (green), bassoon (blue) and CtBP2 (red), imaged in 2D-STED (CtBP2) and confocal mode (homer1 and bassoon). The morphology of afferent synapses of piccolino deficient IHCs appears to be altered with small “compact” ribbons and larger PSDs. Scale bars = 0.5 μm. **(D)** Box plot representing unaltered total number of ribbons (number of CtBP2 puncta; n_IHC_ = 81, n_Corti_ = 8, N_animals_ = 6 for *Pclo^wt/wt^*; and n_IHC_ = 79, n_Corti_ = 7, N_animals_ = 4 for *Pclo^gt/gt^*; *P* = 0.466, t-test) and number of anchored ribbons (number of CtBP2-bassoon juxtapositions; n_IHC_ = 52, n_Corti_ = 6, N_animals_ = 4 for *Pclo^wt/wt^* and n_IHC_ = 51, n_Corti_ = 5, N_animals_ = 3 for *Pclo^gt/gt^*; *P* = 0.681, t-test). Inlaid points represent average numbers/IHC drawn from individual organs of Corti. **(E, F)** Box plots of the 2D area of synaptic ribbons and PSDs respectively, derived by fitting a 2D-Gaussian function to immunofluorescence data as represented in (C). **(E)** n_ribbons_ = 74, N_animals_ = 3 for *Pclo^wt/wt^* and n_ribbons_= 71, N_animals_ = 3 for *Pclo^gt/gt^*; *P* < 0.001, Mann-Whitney-Wilcoxon test. **(F)** n_PSDs_ =70, N_animals_ = 3 for *Pclo^wt/wt^* and n_PSDs_ = 65, N_animals_ = 3 for *Pclo^gt/gt^*; *P* < 0.001, Mann-Whitney-Wilcoxon test. **(G, H)** The distance between the centres of mass of CtBP2 and bassoon spots shows a reduction for piccolino-deficient ribbons (*P* < 0.01, Mann-Whitney-Wilcoxon test), while the distance between the centres of mass of CtBP2 and homer1 was comparable (*P* > 0.05, Mann-Whitney-Wilcoxon test). n_pairs_ =75, N_animals_ = 3 for *Pclo^wt/wt^*; n_pairs_= 69, N_animals_ = 3 for *Pclo^gt/gt^*. Throughout, box and whisker plots present median, lower/upper quartiles and 10–90th percentiles with individual data points overlaid and means shown as crosses.

We then acquired images of *Pclo^wt/wt^* and *Pclo^gt/gt^* synapses using 2D-STED (CtBP2) and confocal (bassoon, homer1) imaging to assess their morphology (Fig 2C). Visually, a large proportion of synaptic ribbons from *Pclo^gt/gt^* rats appeared smaller and more compact, in contrast to the typical ellipsoid-shaped ribbons from *Pclo^wt/wt^* rats. A 2D-Gaussian function was fitted to raw images of randomly selected synapses to estimate the area of the ribbon and the PSD. We found that on average, the size of the synaptic ribbon appeared to be reduced for *Pclo^gt/gt^* rats (38.34x10^3^ ± 2.36x10^3^ nm^2^, S.D. = 19.71x10^3^, n_ribbons_ = 71, N_animals_ = 3) compared to *Pclo^wt/wt^* (46.32x10^3^ ± 1.77x10^3^ nm^2^, S.D. = 15.12x10^3^, n_ribbons_ = 74, N_animals_ = 3; *P* < 0.001, Mann-Whitney-Wilcoxon Test, Fig 2E). Images with more than one ribbon per synapse (observed only in *Pclo^wt/wt^* IHCs) were excluded from analysis to avoid overestimation of ribbon size in *Pclo^wt/wt^* IHCs. On the other hand, the PSD size appeared to be larger for *Pclo^gt/gt^* rats (22.46x10^4^ ± 1.20x10^4^ nm^2^, S.D. = 9.56x10^4^, n_PSD_ = 65, N_animals_ = 3) compared to *Pclo^wt/wt^* rats (17.25x10^4^ ± 1.04x10^4^ nm^2^, S.D. = 8.61x10^4^, n_PSD_ = 70, N_animals_ = 3; *P* < 0.001, Mann-Whitney-Wilcoxon Test, Fig 2F).

Previously, Jing and co-workers have reported ribbon anchorage defects in bassoon gene-trap mutant mice (Jing et al., 2013). To check for any analogous phenotype upon piccolino disruption, we estimated the centers of mass of CtBP2, bassoon and homer1 signals and calculated the ribbon-AZ and the ribbon-PSD center of mass distance. We found a reduction in the ribbon-AZ center of mass distance (n_pairs_ = 69, N_animals_ = 3 for both groups; *P* < 0.01, Mann-Whitney-Wilcoxon Test) in *Pclo^gt/gt^* synapses, whereas the ribbon-PSD center of mass distance appeared comparable (n_pairs_ = 64, N_animals_ = 3 for both groups; *P* > 0.05, t-test) (Fig 2G, H). We speculate that the smaller ribbon size in *Pclo^gt/gt^* IHCs may have resulted in the smaller ribbon-AZ center of mass distance, while this might not have become obvious for ribbon-PSD due to their larger and more variable center of mass distance.

### Large 3D electron microscopic volume reconstructions revealed two distinct ribbon morphologies in mutant IHCs

In a next step, we performed an ultrastructural analysis in order to determine fine structural changes beyond those amenable to light microscopy. Focused ion beam-scanning electron microscopy (FIB-SEM) enabled us to quantify morphological parameters such as ribbon and PSD sizes as well as SV numbers and their localization within IHCs.

We compared the ribbon volume and localization in 2-3 months old adult *Pclo^gt/gt^* and *Pclo^wt/wt^* IHCs (Fig 3A, B; Movie 1,2). While our immunohistochemical data revealed ribbons of *Pclo^gt/gt^* AZs to be smaller on average, FIB-SEM could distinguish two categories of ribbon-type AZs in *Pclo^gt/gt^* IHCs. They could be clearly separated by several morphological features: category 1 encompassed ribbons that appeared comparable to *Pclo^wt/wt^* littermate control ribbons regarding size, shape and SV occupancy. Ribbons sorted in category 2 appeared smaller, more round in shape and lacked SVs at their membrane-distal side (Fig 3C). A first guess that these categories might simply reflect the previously described ribbon number and size difference of modiolar and pillar AZs of IHCs (Hua et al., 2021; Liberman et al., 2011; Michanski et al., 2019; Ohn et al., 2016; Payne et al., 2021), did not match our data. Category 2 ribbons seemed to be equally present at the modiolar and the pillar sides of *Pclo^gt/gt^* IHCs. Due to the low IHC number for the large 3D volume data set we refrained from a statistical analysis. Since the two ribbon categories robustly differ in size and SV numbers, we analyzed all mutant ribbons together and additionally analyzed both categories separately. We found that ribbons of category 2 were indeed smaller with fewer SVs (Fig 3D-F; Appendix Table 1) in contrast to category 1 and *Pclo^wt/wt^* ribbons (2.23x10^6^ nm^3^ vs. 3.07x10^6^ nm^3^ in *Pclo^gt/gt^* category 1 and 6.74x10^6^ nm^3^ in *Pclo^wt/wt^*, i.e., category 2 ribbons being 3 times smaller compared to *Pclo^wt/wt^* ribbons). The PSD area appeared unchanged for all mutant synapses compared to *Pclo^wt/wt^* littermate controls (Fig 3G; Appendix Table 1), which contrasts our immunofluorescence analysis. Notably, no double ribbons per synaptic contact were observed in *Pclo^gt/gt^* rats, while in both FIB-SEM data sets of *Pclo^wt/wt^* IHCs double ribbons appeared frequently (Fig 3C), in accordance with previous observations for mouse IHCs (Hua et al., 2021; Michanski et al., 2019; Payne et al., 2021; Sobkowicz et al., 1982; Sobkowicz et al., 1986; Stamataki et al., 2006; Wong et al., 2014) and our immunofluorescence results.

**Figure 3:**
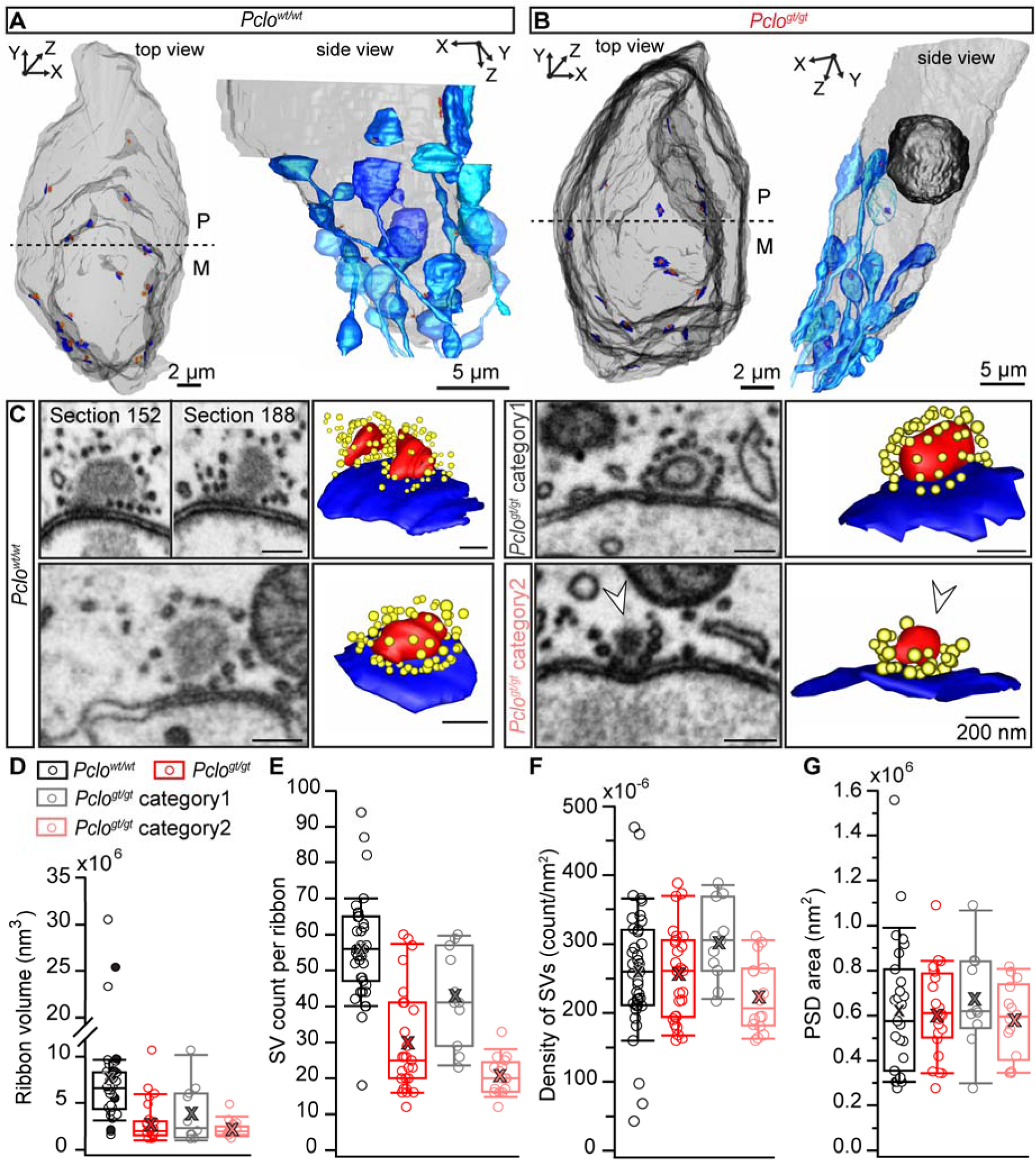
FIB-SEM revealed two morphological distinct ribbon categories in *Pclo^gt/gt^* inner hair cells. **(A, B)** 3D visualizations of *Pclo^wt/wt^* and *Pclo^gt/gt^* (2-3 months old) rat IHCs using FIB-SEM displayed in top view and side view, without or with afferent nerve fiber contacts (different shades of light blue). Based on the tissue context and the cellular apical-basal axis, the separation of pillar (P) and modiolar (M) sides was determined. Light gray: IHC membrane, dark gray: nucleus, red: ribbons, dark blue: postsynaptic density (PSD). **(C)** Representative single sections (left panels) of the reconstructed (right panels) ribbons (red) with their surrounding synaptic vesicles (SVs, yellow) and opposing PSDs (dark blue). Multi-ribbon AZs were observed only in *Pclo^wt/wt^* AZs (left upper panel), while *Pclo^gt/gt^* IHCs revealed two morphological distinct AZ categories (right panel). In contrast to the *Pclo^wt/wt^*-resembling ribbon synapse architecture in category 1, category 2 ribbon synapses take a roundish shape and lack SVs at the upper side of the ribbon (white arrowheads). **(D-G)** Box plots with the mean values (cross) and individual data points display the quantification of ribbon volumes (D), SV counts (E), SV densities (F, SV count values normalized to the ribbon area) and the PSD area (G), for *Pclo^wt/wt^* and *Pclo^gt/gt^* samples, respectively as well as an additional separation of the *Pclo^gt/gt^* data into category 1 and category 2 synapses. Black filled circles in D represent measurements from multi-ribbons AZs. N_animals_ = 2, n_IHCs_ = 2 respectively for *Pclo^wt/wt^* and *Pclo^gt/gt^* condition.

**Appendix Table 1:** FIB-SEM data on count, size and distance measurements of AZ parameters. Data are presented as mean ± SEM. Abbreviations: SV = synaptic vesicles, PSD = postsynaptic density, NND = nearest neighbor distance.

|  | *Pclo*<sup>wt/wt</sup> | *Pclo*<sup>gt/gt</sup> | *Pclo*<sup>gt/gt</sup> category 1 | *Pclo*<sup>gt/gt</sup> category 2 |
| --- | --- | --- | --- | --- |
| **Ribbon volume (nm<sup>3</sup>) x10<sup>6</sup>** | 7.82 ± 0.95 | 3.09 ± 0.42 | 4.00 ± 0.92 | 2.37 ± 0.21 |
| **SV count** | 56.05 ± 2.22 | 29.78 ± 2.76 | 42.91 ± 3.99 | 20.75 ± 1.31 |
| **SV density x10<sup>-6</sup>** | 259.96 ± 14.29 | 261.05 ± 12.76 | 305.91 ± 17.18 | 206.32 ± 13.03 |
| **PSD area (nm<sup>2</sup>) x10<sup>6</sup>** | 0.63 ± 0.06 | 0.60 ± 0.04 | 0.68 ± 0.07 | 0.58 ± 0.04 |
| **NND mitochondria (nm)** | 198.04 ± 27.13 | 185.16 ± 31.27 | 234.79 ± 52.74 | 160.34 ± 38.29 |
| **NND ribbons (μm)** | 3.40 ± 0.38 | 4.08 ± 0.58 | 2.89 ± 0.26 | 4.77 ± 0.85 |

Next to ribbon morphology, the overall distribution of ribbons within an IHC might be changed in the mutants. Therefore, we quantified the nearest neighbor distance between ribbons in both genotypes in our FIB-SEM data (Fig EV1E). For *Pclo^wt/wt^* IHCs, we included the distance measurements between ribbons of the same synaptic contact (indicated as filled points in Fig EV1E). However, no differences between all *Pclo^gt/gt^* compared to *Pclo^wt/wt^* ribbons could be observed, suggesting that the distribution of synaptic contacts is normal in *Pclo^gt/gt^* IHCs (Fig EV1E; Appendix Table 1). An analogous analysis of nearest neighbor distances between ribbons from our immunolabeled confocal sections of the organs of Corti confirmed these findings since no difference in distances could be found (Fig EV1G, H). We further noted that mitochondria often appeared in close vicinity to category 2 mutant ribbon synapses (Fig EV1A-C). Indeed, apposition of mitochondria to category 2 ribbons was closer compared to *Pclo^wt/wt^* and *Pclo^gt/gt^* category 1 ribbons (Fig EV1D; Appendix Table 1). However, the overall frequency of mitochondria being in close vicinity to ribbons was comparable between *Pclo^gt/gt^* and *Pclo^wt/wt^* IHCs (Fig EV1C). Finally, FIB-SEM revealed that larger ribbons frequently harbor a translucent core, regardless of the genotype (Fig 3C, Fig EV1F), which is in line with previous findings (Liberman, 1980; Michanski et al., 2019; Sobkowicz et al., 1982; Stamataki et al., 2006). Interestingly, no translucent core could be found in the category 2 ribbons of *Pclo^gt/gt^* rats, likely owing to their smaller size.

**Fig EV1:**
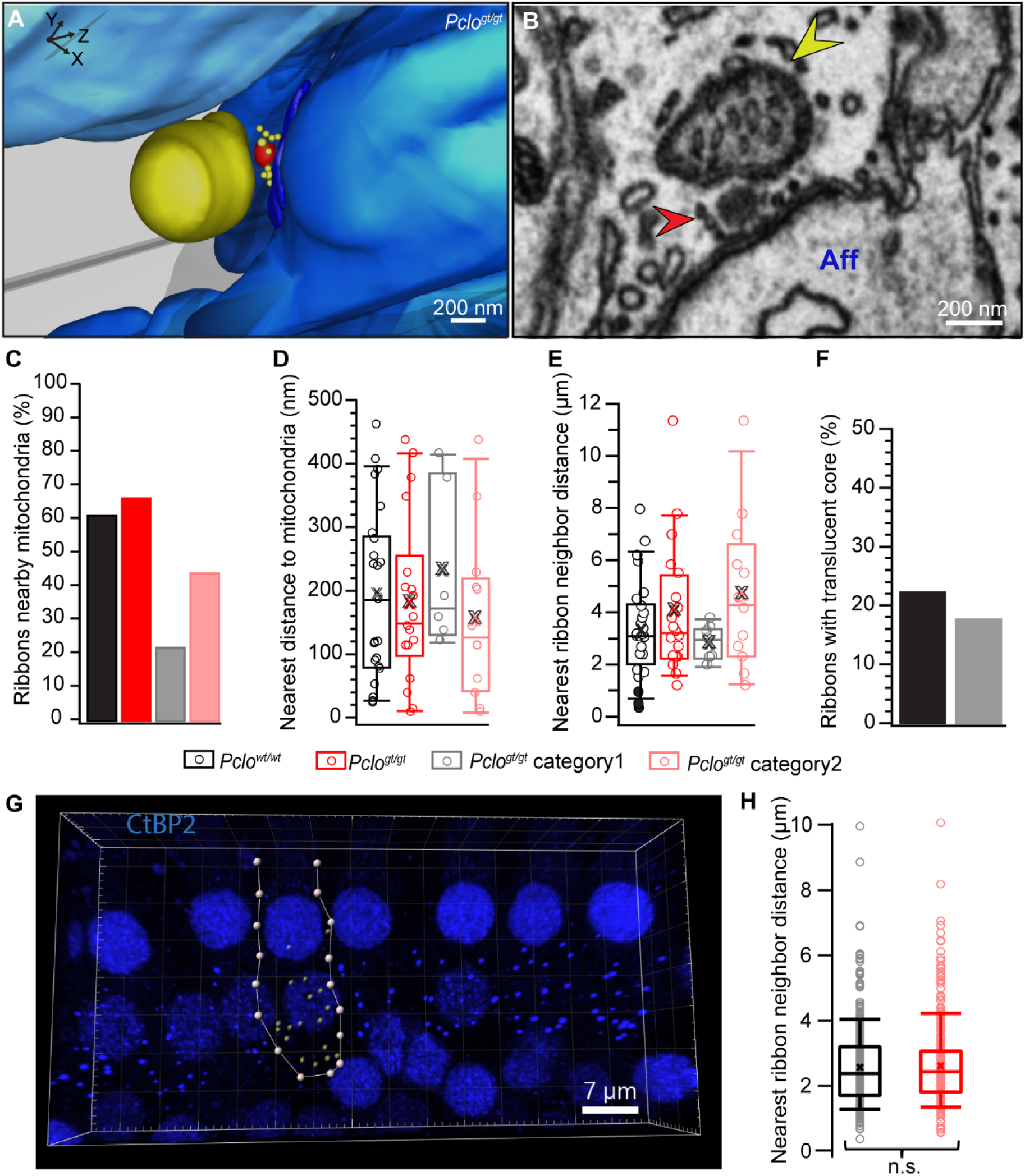
Mitochondria tend to be closer to the ribbons of category 2 *Pclo^gt/gt^* AZs. **(A)** FIB-SEM 3D model and an exemplary section **(B)** of a representative *Pclo^gt/gt^* category 2 ribbon (red/arrowhead) with a mitochondrion (dark yellow/arrowhead) nearly touching the upper side of the ribbon (devoid of SVs) with an afferent bouton (Aff). **(C)** Analysis of mitochondria-proximal ribbons in a range of 500 nm around the ribbon. **(D)** Measurements of the shortest distance between synaptic ribbon and mitochondria surfaces. **(E)** Box plot of distance measurements, measured from the surfaces of the respective structure, between nearest neighboring synaptic ribbons depicted with individual data points and mean values highlighted as a cross. Black filled circles represent the distance measurements between the individual ribbons of multi-ribbon AZs. **(F)** Quantification of ribbon counts exhibiting a translucent core reveals this feature only in *Pclo^wt/wt^* ribbons and the larger sized category 1 *Pclo^gt/gt^* ribbons (gray) but not in category 2 *Pclo^gt/gt^* ribbons. N_animals_ = 2, n_IHCs_ = 2 respectively for *Pclo^wt/wt^* and *Pclo^gt/gt^* condition. **(G)** Immunofluorescent spots corresponding to synaptic ribbons (labeled with an antibody against CtBP2/RIBEYE) were detected using Imaris 9.6 (Oxford Instruments) in confocal 3D projections of apical organs of Corti from 2 months old *Pclo^wt/wt^* and *Pclo^gt/gt^* rats. Yellow spots in exemplary image shown here represent detected ribbons within one IHC. **(H)** Nearest ribbon neighbor distance was measured between spot centers and appears comparable between *Pclo^wt/wt^* and *Pclo^gt/gt^* conditions, (*P* > 0.05, Mann-Whitney-Wilcoxon test), in agreement with FIB-SEM data shown in (E). n_ribbons_ = 435, n_IHCs_ = 33, N_animals_ = 3 for *Pclo^wt/wt^* and n_ribbons_ = 396, n_IHCs_ = 32, N_animals_ = 3 for *Pclo^gt/gt^*.

Transmission electron microscopy of ultrathin sections from conventional embeddings further corroborated these findings: Translucent cores are present in larger ribbons, while smaller ribbons in *Pclo^gt/gt^* IHCs showed a uniform electron-density and lacked SVs at the upper ribbon part (Fig EV2A,B) and were thus characterized as category 2. Using an anti-piccolino immunogold labeling with the same antibody as for our immunofluorescence analysis, we found that piccolino was almost equally distributed along individual *Pclo^wt/wt^* ribbons of 10-11 months of age, while it seems that ribbons are less frequently labeled in *Pclo^wt/gt^* IHCs suggesting a reduction of piccolino molecules in heterozygotes (Fig EV2C,D).

#### Movie 1 and 2: FIB-SEM visualizations of the nuclear and basal region of *Pclo^wt/wt^* and *Pclo^gt/gt^* cochlear IHCs with corresponding 3D segmentations

Movies scanning through the FIB-SEM z-stacks of *Pclo^wt/wt^* (Movie 1, 2 months and 4 days old) and *Pclo^gt/gt^* (Movie 2, 3 months old) IHCs. The displayed 3D models depict IHC contours (transparent gray), part of the nuclei (dark gray), innervating afferent nerve fibers (blue), ribbon synapses (red), their corresponding SVs (yellow) and PSDs (dark blue). While multiribbon AZs (highlighted with red arrow) can be detected in *Pclo^wt/wt^* IHCs, exclusively single ribbons that can be divided into two morphological distinct categories (highlighted with white arrows) are found in *Pclo^gt/gt^* IHCs.

Movie1: https://owncloud.gwdg.de/index.php/s/lLuJangpAovxArc

Movie2: https://owncloud.gwdg.de/index.php/s/XFPfoGcwWaMH4H3

**Fig EV2:**
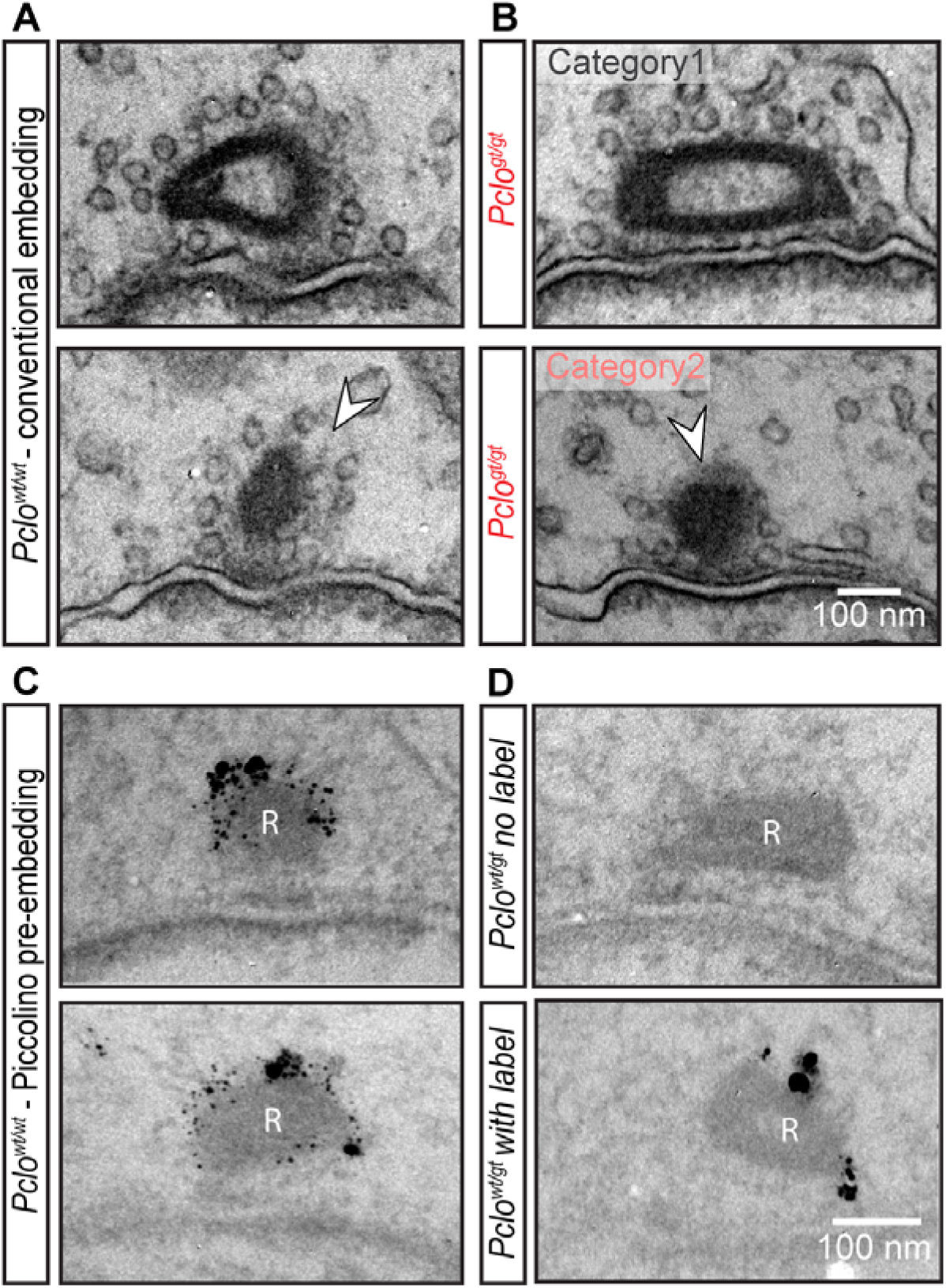
2D Transmission electron microscopy corroborates the notion of 2 ribbon categories in *Pclo^gt/gt^* as well as in *Pclo^wt/gt^* inner hair cells – immuno-electron microscopical detection of piccolino. **(A)** *Pclo^wt/wt^* ribbons with (upper panel) and without a translucent core (lower panel). Ribbons show a SV distribution along the full ribbon surface (arrowhead). **(B)** *Pclo^gt/gt^* category 1 (upper panel) with a translucent core and category 2 (lower panel) with a lack of SVs at the upper ribbon side (arrowhead). **(C, D)** Representative examples of an anti-piccolino pre-embedding immunogold labeling, which can be found around the synaptic ribbons (R) in *Pclo^wt/wt^* (C) and *Pclo^wt/gt^* (D) IHCs. (D) In *Pclo^wt/gt^* IHCs, most synaptic ribbons lacked anti-piccolino labeling (upper panel). However, for few ribbons the occurrence of silver enhanced gold particles could be observed (lower panel).

### Electron tomography reveals fewer ribbon-associated SVs at category 2 mutant AZs but a normal fraction of ribbon-SV filaments

Next, we analyzed the morphologically distinct SV pools of ribbon synapses and SV tethering as piccolino was shown to interact with RIBEYE and proposed to support the SV trafficking via interacting with other presynaptic proteins. We studied ribbon-associated (RA) and membrane-proximal (MP) SV pools, as well as their tethering using electron tomography combined with high-pressure freezing and freeze substitution (HPF/FS) which enables high resolution with a close-to-native structural preservation of ribbons, SVs and filaments (Chakrabarti et al., 2018).

We could verify the two ribbon categories in *Pclo^gt/gt^* IHCs, with normal *Pclo^wt/wt^* resembling ribbons (category 1) and altered, small and spherical ribbons, void of membrane-distal SVs (category 2) (Fig 4A-C). Consequently, as for the FIB-SEM data, we determined all morphometric parameters (shown in Fig 4D) for all *Pclo^gt/gt^* AZs combined and separated into category 1 and 2. For the grouping of the reconstructed ribbons of *Pclo^gt/gt^* rats, we performed the K-means clustering analysis, which confirmed our blinded and manual annotation of AZs into category 1 and 2, including all *Pclo^gt/gt^* and *Pclo^wt/wt^* tomograms. Notably, in the manual analysis, none of the *Pclo^wt/wt^* AZs were sorted in category 2, confirming that this is a morphological category only found in the mutants. Considering all mutant ribbons (*Pclo^gt/gt^*), most of the parameters were comparable to *Pclo^wt/wt^* (Fig 4E-G; Appendix Table 2), but significantly fewer SVs were observed at *Pclo^gt/gt^* AZs (Fig 4H), while the differences in SV density did not reach significance (Fig 4I). Furthermore, we observed a consistent difference in all our morphometric parameters between category 1 and 2 *Pclo^gt/gt^* AZs. Interestingly, the ribbon volume, the PSD area, the total SV count per ribbon as well as the RA-SV count were even increased in category 1 AZs in comparison to *Pclo^wt/wt^* AZs (Fig 4E,G,H,K; Appendix Table 2). However, the PSD area difference did not reach significance in contrast to our immunofluorescence data. This discrepancy might arise from the limited section thickness of 250 nm for electron tomography, which does not allow for a full reconstruction of the PSD extent.

**Figure 4:**
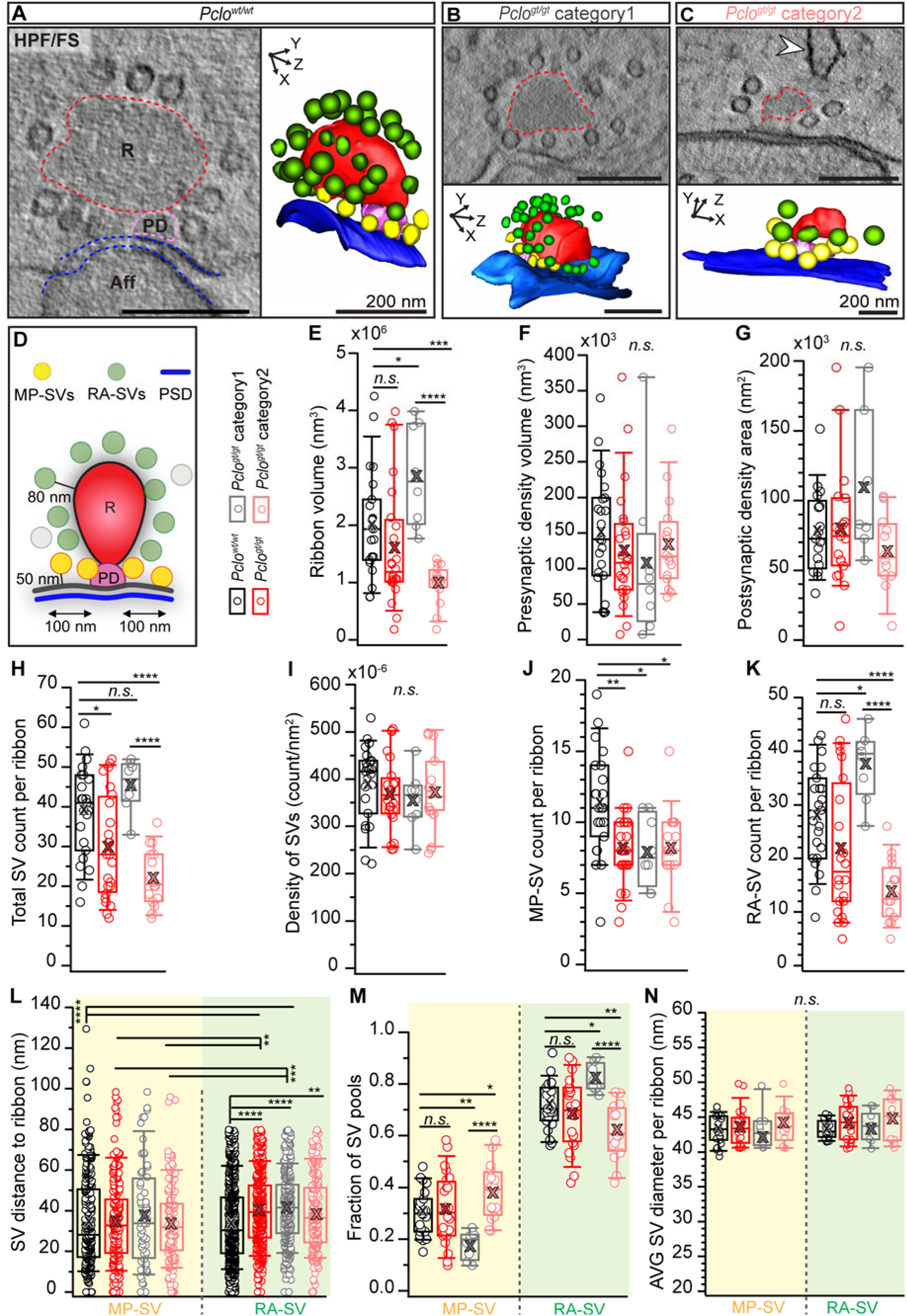
Electron tomography reveals smaller ribbons and fewer RA-SVs in *Pclo^gt/gt^* category 2 AZs. **(A-C)** Representative virtual electron tomographic sections with their corresponding 3D models of *Pclo^wt/wt^* (A,) and *Pclo^gt/gt^* IHCs (B, C) obtained after HPF/FS from 1-2 months old rats. Reconstructions (right panels) show the ribbon (R in left panels) in red, the presynaptic density (PD) in magenta, membrane-proximal synaptic vesicles (MP-SVs) in yellow, ribbon-associated synaptic vesicles (RA-SVs) in green, the postsynaptic density (PSD) of the afferent bouton terminal (Aff) in blue. Based on an unsupervised K-means clustering, *Pclo^gt/gt^* ribbon synapses were also divided into category 1 and category 2 AZs. Compared to *Pclo^wt/wt^* and category 1 AZs, the upper (membrane-distal) ribbon side of category 2 AZs was devoid of RA-SVs but frequently faced cisternal structures (arrowhead). **(D)** Schematic illustration of tomogram analysis. **(E-N)** Box plots with individual data points and mean values highlighted as crosses show the quantification of the specific parameters. While smaller ribbons were observed for *Pclo^gt/gt^* category 2 AZs and larger sized ribbons for *Pclo^gt/gt^* category 1 (E), no significant differences were detected for PD (F) and PSD (G) sizes. Significantly less SVs per ribbon were observed in *Pclo^gt/gt^* deriving mainly from the category 2 (H). The SV density (SV counts normalized to the ribbon area) was unaltered in *Pclo^gt/gt^* AZs (pooled or per category). Analysis of the two different SV pools showed equally reduced MP-SV counts for all *Pclo^gt/gt^* analysis groups (J). Significantly fewer RA-SVs were found in the *Pclo^gt/gt^* category 2, while more were present in *Pclo^gt/gt^* category 1 (K). Nearest distance measurements from the SV to the ribbon demonstrated increased distances for the RA-SV pool in all three *Pclo^gt/gt^* analysis groups (L). Opposing results regarding the SV pool fraction between category 1 and category 2 *Pclo^gt/gt^* AZs were detected (M). While a greater fraction of MP-SVs was found in the category 2 than in category 1, the RA-SV fraction was significantly decreased in category 2 in comparison to category 1. **(N)** No differences between *Pclo^wt/wt^* and *Pclo^gt/gt^* in the SV diameter were observed for either SV pool. *Pclo^wt/wt^*: N_animals_ = 3, n_ribbons_ = 23, n_PSDs_ = 17; *Pclo^gt/gt^*: N_animals_ = 4, n_ribbons_ = 24, n_PSDs_ = 19; Category 1: n_ribbons_ = 8, n_PSDs_ = 7; Category 2: n_ribbons_ = 16, n_PSDs_ = 12. Significant differences between two groups were analysed with the t-test or the Mann–Whitney Wilcoxon test (E-K, M). For multiple comparisons, ANOVA followed by the post-hoc Tukey (E-K, M, N) or KW test followed by NPMC test (L) was performed. For more detailed information see also Appendix Table 2, 3.

**Appendix Table 2:**
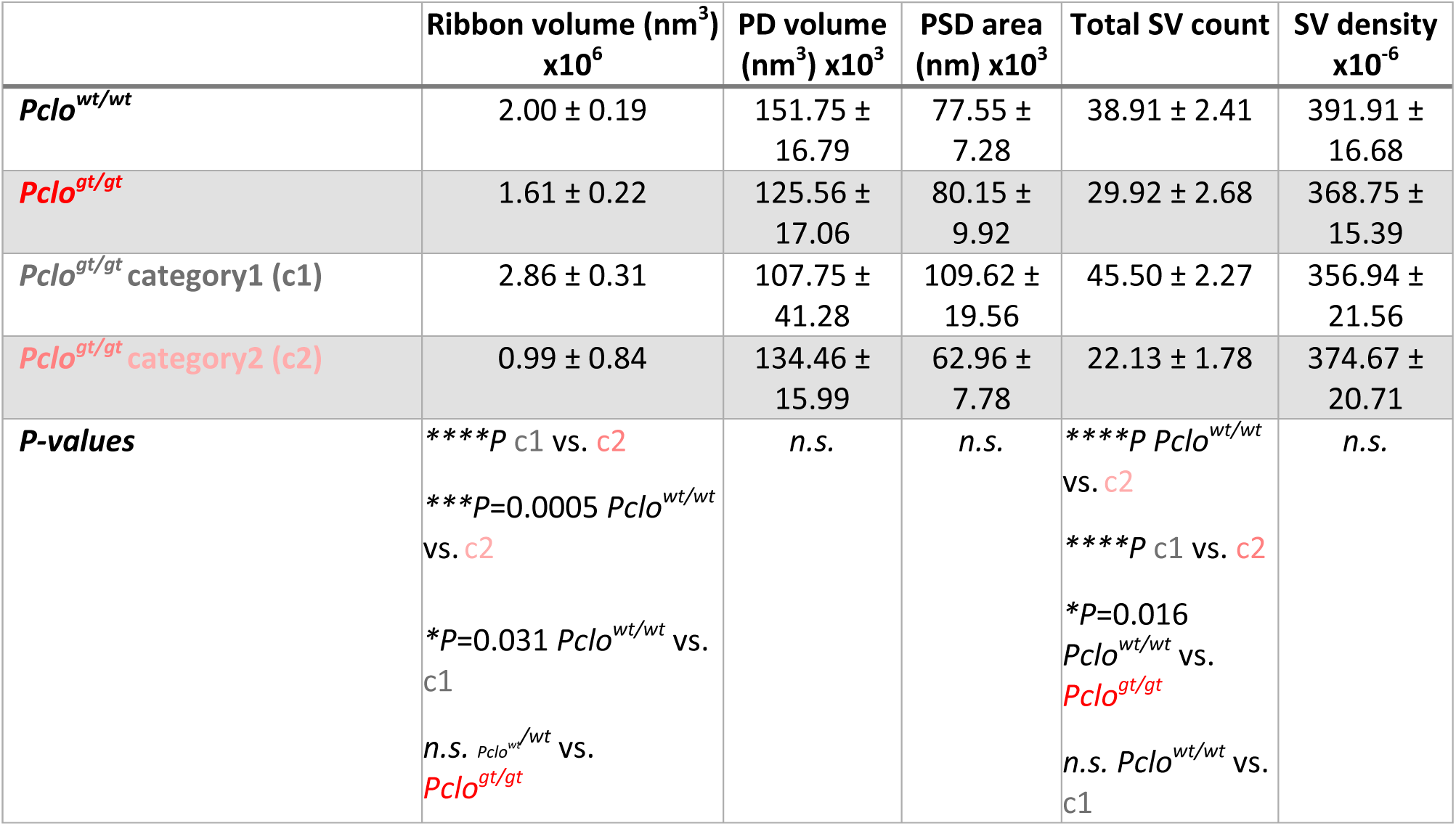
Counts and size measurements of AZ structures observed by HPF/FS electron tomography. Data are presented as mean ± SEM. *P*-values between two groups were calculated using the t-test or the Mann–Whitney Wilcoxon test. For multiple comparisons, the ANOVA test with post-hoc Tukey test or KW followed by NPMC test was performed. Non-significant differences are indicated as *n.s.*, significant differences as *\*P* < 0.05, *\*\*P* < 0.01, *\*\*\*P* < 0.001, *\*\*\*\*P* < 0.0001. Abbreviations: PD = presynaptic density, PSD = postsynaptic density, SV = synaptic vesicle.

Ultrastructural quantification of the two morphological SV pools (described in (Chakrabarti et al., 2018)) revealed a reduction in the number of MP-SVs in category 1 and 2 AZs separately as well as in the combined mutant dataset (Fig 4J; Appendix Table 3). Consistent with our FIB-SEM data, ribbons of category 2 AZs lacked SVs at their membrane-distal side (Fig 4K). However, this did not affect the SV density per ribbon surface area (Fig 4I). Moreover, RA-SV seemed to be further away from the synaptic ribbon in both mutant AZ categories (Fig 4L; Appendix Table 3). Next, we investigated the fractions of the SV-pools following our previously published approach (Chakrabarti et al., 2018). The MP-and the RA-SV pool fractions were comparable between the combined mutant ribbon synapses and the littermate controls (Fig 4M; Appendix Table 3). As expected from the above results, category 1 and category 2 AZs showed opposing effects: while in category 1, the fraction of MP-SVs was significantly reduced, it was strikingly increased in *Pclo^gt/gt^* category 2 AZs, which was significant when compared to *Pclo^gt/gt^* category 1 and *Pclo^wt/wt^* AZs. Such an opposite effect was also found for the RA-SVs that were fewer in category 2 but increased in category 1 compared to *Pclo^wt/wt^* AZs, explaining why we did not observe overall changes when combining all mutant AZs (Fig 4M; Appendix Table 3). The analysis of SV sizes from the different pools resulted in uniformly sized diameters for all groups (Fig 4N; Appendix Table 3).

**Appendix Table 3:**
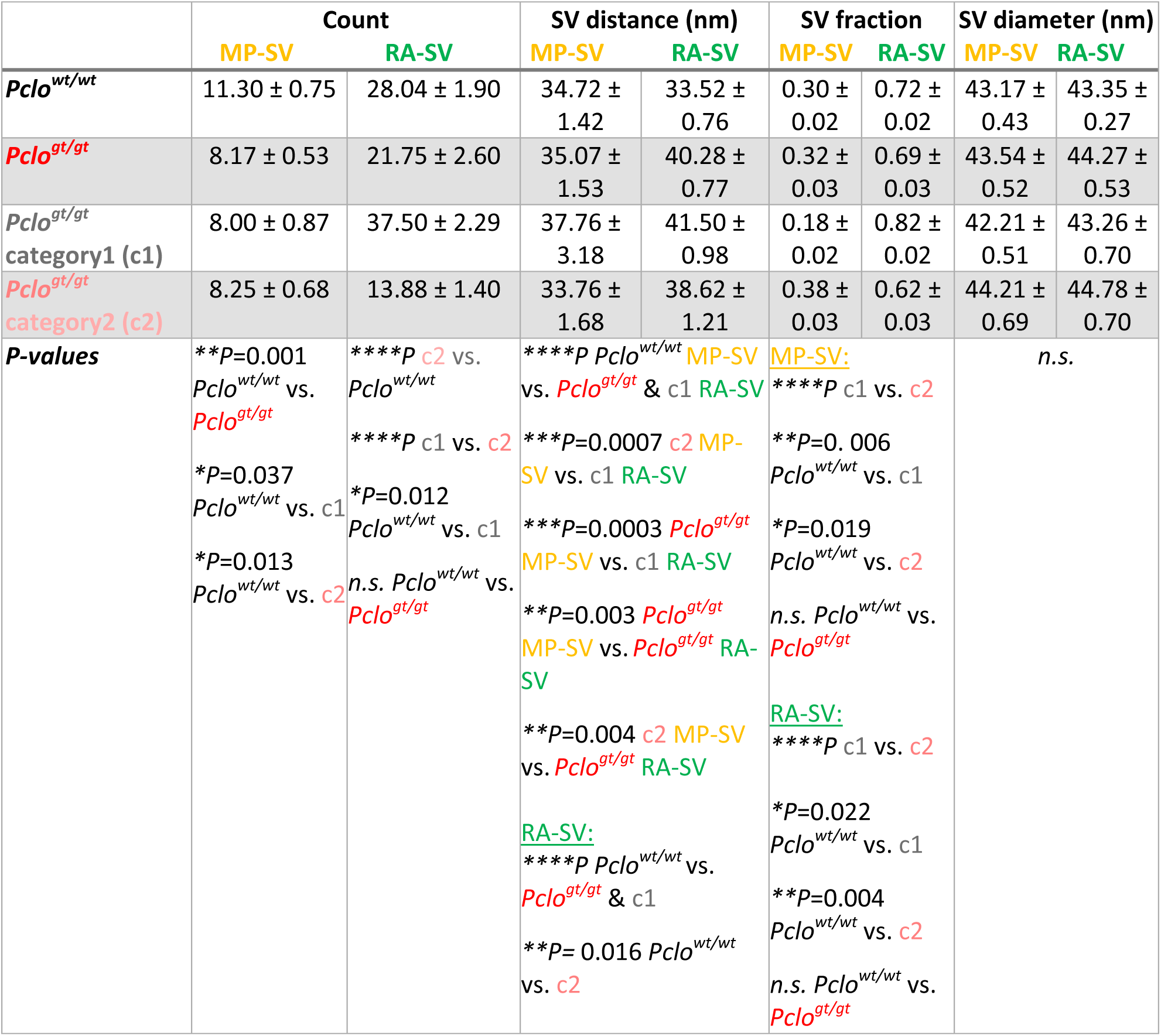
Counts, size and distance measurements of MP and RA-SV pools from HPF/FS electron tomography. Data are presented as mean ± SEM. Depending on the normality and equality of variances, significant differences between two groups were analyzed with the t-test or the Mann–Whitney Wilcoxon test. For multiple comparisons, the ANOVA test with post-hoc Tukey test or KW followed by NPMC test was performed. Non-significant differences are indicated as *n.s.*, significant differences as *\*P* < 0.05, *\*\*P* < 0.01, *\*\*\*P* < 0.001, *\*\*\*\*P* < 0.0001. Abbreviations: MP-SV = membrane-proximal synaptic vesicle, RA-SV = ribbon-associated synaptic vesicle.

As tethering of SVs to the ribbon might involve ribbon-standing piccolino, we investigated the tethering of the RA-SVs analyzing the filament number and to which structures the SVs connect to. These different tethering states were previously stated as morphological SV subpools, which are present at the ribbon as well as at the membrane (Chakrabarti et al., 2018). In this study, we focused on the RA-SVs since piccolino was shown to localize exclusively to the ribbon (Dick et al., 2001, 20; Limbach et al., 2011; Michanski et al., 2019). Filaments could be observed at both, *Pclo^wt/wt^* and *Pclo^gt/gt^* ribbon synapses (Fig 5A,B). When counting the number of filaments per ribbon (Fig 5D) or alternatively the numbers of all filaments per tomogram (Fig 5E), we did not observe any significant differences (Appendix Table 4). Subsequently, we quantified the different RA-SV-subpools as done in (Chakrabarti et al., 2018) but no changes could be detected (Fig 5F; Appendix Table 4). Therefore, we conclude that tethering of the remaining ribbon-associated SVs of piccolino-deficient IHC synapses is unchanged.

**Figure 5:**
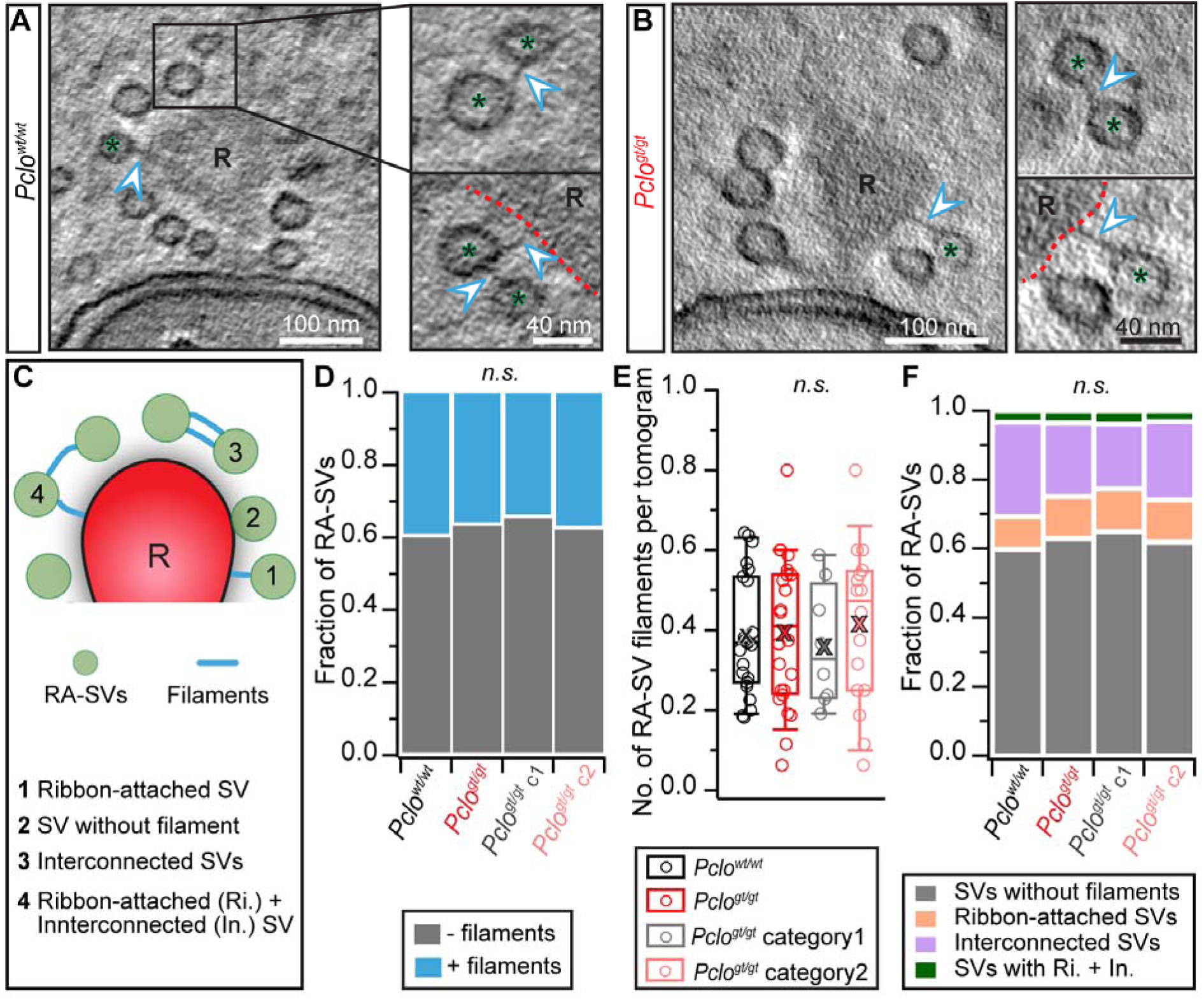
Normal tethering of synaptic vesicles at *Pclo^gt/gt^* ribbons. **(A,B)** Exemplary virtual electron tomographic sections of *Pclo^wt/wt^* and *Pclo^gt/gt^* ribbon synapses (ribbon highlighted with “R” and/or red dotted line). Proteinaceous filaments of different types (Ri: ribbon-attached, In: interconnected, and both Ri and In; filaments are marked with arrowheads) were detected at RA-SVs (asterisks) in both *Pclo^wt/wt^* and *Pclo^gt/gt^* IHCs. **(C)** Illustration of the various tethering states of RA-SVs. **(D,E)** The fraction of RA-SVs with and without filaments (D) as well as the total number of RA-SV filaments per tomogram (E) in *Pclo^gt/gt^* was comparable to *Pclo^wt/wt^* (t-test and ANOVA with post-hoc Tukey test). **(F)** The graph represents the fraction of RA-SVs around the ribbon regarding the filament type. No changes in the occurrence of individual filament types could be found. Multiple comparison tests of the filament types were performed using ANOVA with the post-hoc Tukey test (without filaments, Ri., In.) or KW test followed by NPMC test (Ri. + In.). *Pclo^wt/wt^*: N_animals_ = 3, n_ribbons_ = 23; *Pclo^gt/gt^*: N_animals_ = 4, n_ribbons_ = 24; Category 1: n_ribbons_ = 8; Category 2: n_ribbons_ = 16. For more detailed information see Appendix Table 4.

**Appendix Table 4:** Electron tomographical tether analysis for RA-SVs. Data are presented as mean ± SEM. Depending on the normality and equality of variances tests, significant differences between two groups were analyzed with the t-test or the Mann–Whitney Wilcoxon test. For multiple comparisons, the ANOVA test followed by post-hoc Tukey test or the KW followed by NPMC test was performed. Non-significant differences are indicated as *n.s.*, significant differences as *\*P* < 0.05, *\*\*P* < 0.01, *\*\*P* < 0.001, *\*\*\*\*P* < 0.0001. The abbreviation - stands for without, + for with, Ri. for ribbon attached, In. for interconnected.

|  | Fraction of tethers |  | No. of tethers per tomogram | Fraction of filament types |  |  |  | Distance cisternal structures |
| --- | --- | --- | --- | --- | --- | --- | --- | --- |
|  | - | + |  | - | Ri. | In. | Ri.+In. |  |
| *Pclo*<sup>wt/wt</sup> | 0.60 | 0.40 | 0.39 ± 0.03 | 0.60 | 0.10 | 0.27 | 0.03 | 266.54 ± 22.25 |
| *Pclo*<sup>gt/gt</sup> | 0.63 | 0.37 | 0.40 ± 0.04 | 0.63 | 0.12 | 0.21 | 0.03 | 191.82 ± 22.15 |
| *Pclo*<sup>gt/gt</sup> category1 (c1) | 0.65 | 0.35 | 0.36 ± 0.05 | 0.65 | 0.12 | 0.19 | 0.04 | 179.70 ± 37.24 |
| *Pclo*<sup>gt/gt</sup> category2 (c2) | 0.62 | 0.38 | 0.41 ± 0.05 | 0.62 | 0.12 | 0.23 | 0.03 | 198.10 ± 27.96 |
| **P-values** | *n.s.* |  | *n.s.* | *n.s.* |  |  |  | **\*\*P=0.009**<br>*Pclo*<sup>wt/wt</sup> vs.<br>*Pclo*<sup>gt/gt</sup> |

Finally, we found cisternal structures in close ribbon proximity at *Pclo^gt/gt^* ribbon synapses (Fig. 4C, Fig EV3; Appendix Table 4). In category 2 *Pclo^gt/gt^* IHCs, these cisternal structures could frequently be observed close to the SV lacking upper ribbon part (Fig EV3B). Similar observations were made in IHCs upon disruption of bassoon (Khimich et al., 2005) and the endocytic proteins AP2 and endophilins (Jung et al., 2015b; Kroll et al., 2019). Moreover, similar cisternal structures have been reported in hippocampal synapses of *Pclo^gt/gt^* rats. The precise identity of these structures, which have been referred to as endosome-like vacuoles remains to be determined.

**Fig EV3:**
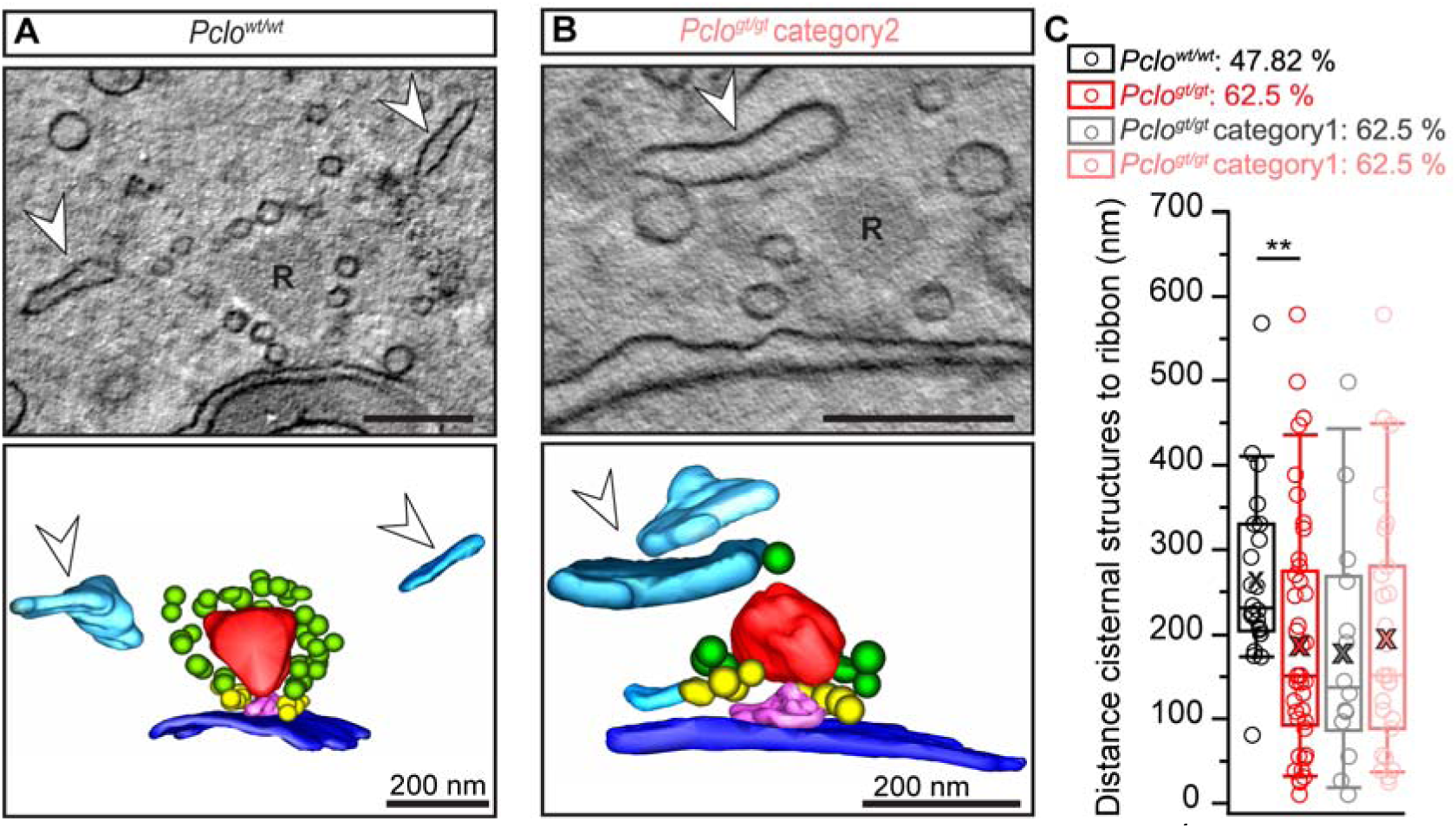
Cisternal structures close to ribbon synapses frequently occur in *Pclo^gt/gt^* IHCs. **(A,B)** Individual virtual electron tomographic sections of HPF/FS IHCs depicting a *Pclo^wt/wt^* (A) and a *Pclo^gt/gt^* (B) ribbon synapse (upper panel) with their corresponding 3D models (lower panel). Ribbons (R) are represented in red, presynaptic densities in magenta, MP-SVs in yellow, RA-SVs in green, postsynaptic densities in dark blue and cisternal structures in cyan, which are additionally highlighted with arrowheads. **(C)** Cisternal structures within a 1 µm range are closer to ribbons at *Pclo^gt/gt^* AZs (*P* = 0.009, Mann–Whitney Wilcoxon test, for multiple comparison KW followed by NPMC test). *Pclo^wt/wt^*: N_animals_ = 3, n_ribbons_ = 23; *Pclo^gt/gt^*: N_animals_ = 4, n_ribbons_ = 24; Category 1: n_ribbons_ = 8; Category 2: n_ribbons_ = 16. For more detailed information see Appendix Table 4.

### Impaired Ca^2+^ channel clustering in mutant animals

Previous studies often reported altered Ca^2+^ channel clustering upon the disruption of presynaptic scaffold proteins at ribbon-type AZs (reviews in Moser et al., 2019; Pangrsic et al., 2018). For instance, disruption of bassoon, RIM, RIM-BP2 as well as RIBEYE impairs Ca^2+^ channel clustering at IHC AZs (Frank et al., 2010; Jean et al., 2018; Jung et al., 2015a; Krinner et al., 2017; Neef et al., 2018). To investigate the impact of piccolino disruption on Ca^2+^ channel clustering, we stained acutely dissected organs of Corti from 2-3 weeks old rats, as employed for IHC physiology (see below), using antibodies against Ca_V_1.3 channels, bassoon and CtBP2 and performed 2D-STED imaging (Fig 6A,B). We randomly selected and imaged 176 synapses from *Pclo^wt/wt^* (N_animals_ = 3) and 191 synapses from *Pclo^gt/gt^* (N_animals_ = 3) and categorized them based on the morphology of Ca^2+^ channel clusters into single-line, double-line, spot-like, and complex clusters (Fig 6B), as has been previously described (Jean et al., 2018; Krinner et al., 2017; Neef et al., 2018). The subjective categorization generated comparable outcomes when performed by different observers. In control littermates, up to 70.5% of Ca_V_1.3 clusters appeared as a single line, while smaller fractions appeared as double lines (10.8%) and spot-like (round) clusters (18.7%). Of the *Pclo^gt/gt^* Ca_V_1.3 clusters, on the contrary, only about 47.7% appeared as single lines and 10.5% as double lines, while 28.8% of them showed a spot-like shape. Additionally, several Ca_V_1.3 clusters in *Pclo^gt/gt^* synapses appeared as fragmented lines and rings seemingly composed of multiple smaller substructures, which we categorized as complex clusters (remaining 13.1%; Fig 6C). Likely as a corollary, we noted more Ca_V_1.3 structures at *Pclo^gt/gt^* AZs than in wild-type (Fig 6D). We cannot rule out that an artificial aggregation of untethered Ca^2+^ channels by the fixative or antibodies contributes to the Ca_V_1.3 clusters remaining as fragments and single spot-like Ca_V_1.3 clusters (Neef et al., 2018).

**Figure 6.**
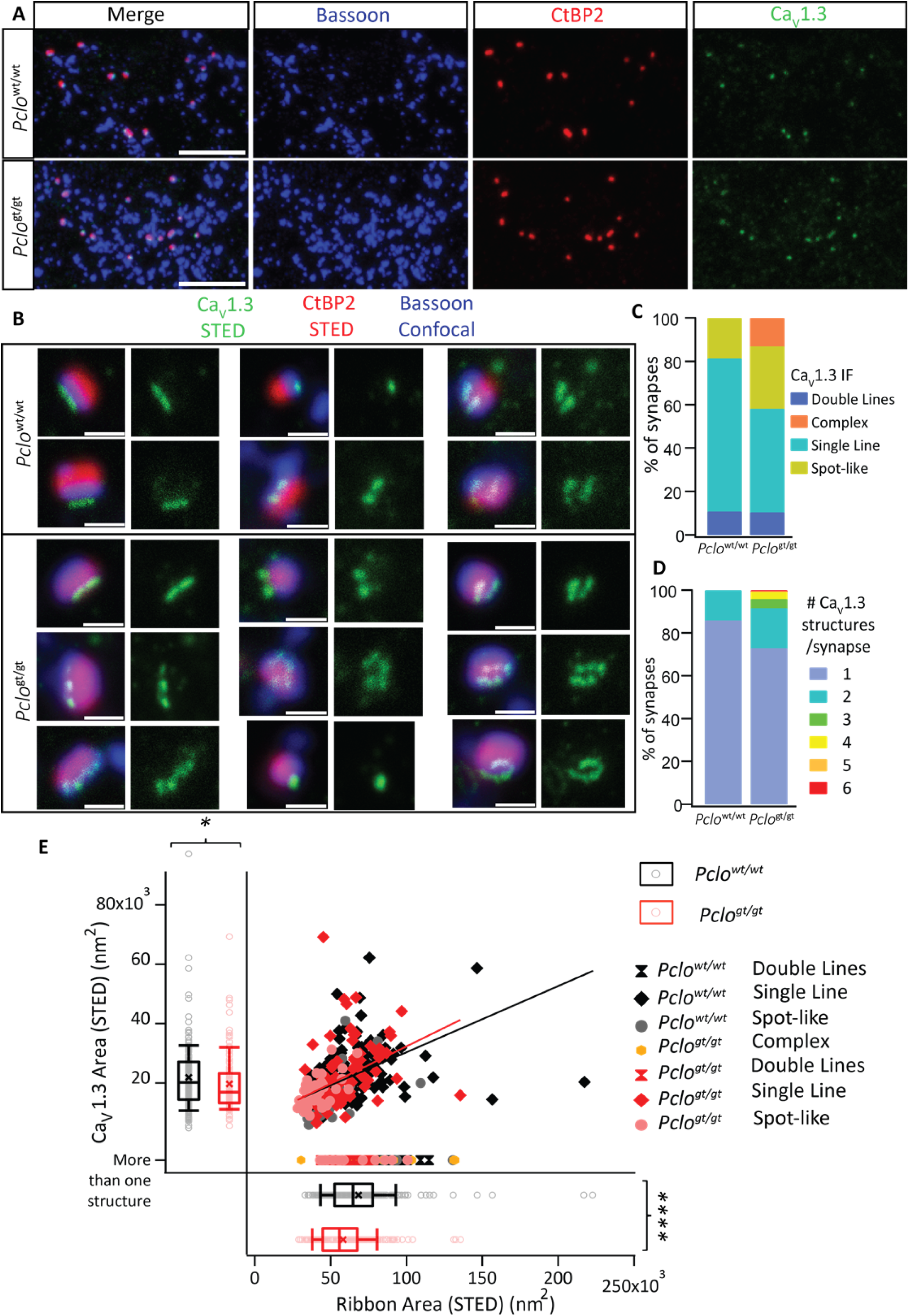
STED imaging reveals altered morphology of Ca_V_1.3 channel clusters upon piccolino disruption. **(A)** Maximum projections of confocal sections of organ of Corti from 2-3 weeks old rats, stained for the presynaptic scaffold protein bassoon (blue), CtBP2 labeling the synaptic ribbon (red) and Ca_V_1.3 channel clusters (green). Juxtaposition of the three proteins appears comparable. Scale bars = 5 µm. **(B)** Randomly selected synapses from the triple-labeling represented in (A) were further imaged using 2D-STED (Ca_V_1.3 and CtBP2) and confocal (bassoon) modes. Scale bars = 0.5 µm. Most synapses in *Pclo^wt/wt^* IHCs showed a single line-shaped Ca_V_1.3 channel cluster, while some showed spot-like clusters and double lines (top panel). On the contrary, *Pclo^gt/gt^* synapses additionally showed Ca_V_1.3 clusters in complex arrangements, with some clusters seemingly broken into several smaller spots (lower panel). **(C, D)** 176 *Pclo^wt/wt^* (N_animals_ = 3) and 191 *Pclo^gt/gt^* (N_animals_ = 3) were categorized based on the apparent morphology of their Ca_V_1.3 channel clusters into line-, double line-, spot-like and complex-clusters. *Pclo^gt/gt^* IHCs evidently show a higher percentage of synapses with spot-like morphology and complex clusters with multiple substructures. **(E)** The 2D areas of the synaptic ribbon and of corresponding Ca_V_1.3 clusters (only ones with a single structure) were estimated by fitting a 2D-Gaussian function to the STED data obtained. The ribbons in IHCs of 2-3 weeks old *Pclo^gt/gt^* rats appear significantly smaller (*P* < 0.0001, Mann-Whitney-Wilcoxon test), even after excluding the outliers in the *Pclo^wt/wt^* dataset. The area of Ca_V_1.3 clusters with single structures (single line/spot; 151 *Pclo^wt/wt^* synapses, 129 *Pclo^gt/gt^* synapses) also appears smaller (*P* < 0.05, Mann-Whitney-Wilcoxon test). Note the positive correlation between ribbon and size of Ca_V_1.3 clusters (P_r_ = 0.53, *P* < 0.0001 for *Pclo^wt/wt^* and P_r_ = 0.44, *P* < 0.0001 for *Pclo^gt/gt^*). Ribbon areas corresponding to clusters with multiple structures have also been plotted. Box plots depict individual data points overlaid with mean values shown as crosses.

Similar to our observations in 2 months old rats, we saw a clear reduction in the size of the synaptic ribbon in younger *Pclo^gt/gt^* rats; 57.65x10^3^ ± 1.36x10^3^ nm^2^, S.D. = 18.75x10^3^, n_ribbons_ = 191, N_animals_ = 3 for *Pclo^gt/gt^ vs.* 67.81x10^3^ ± 1.98x10^3^ nm^2^, S.D. = 26.26x10^3^, n_ribbons_ = 176, N_animals_ = 3 for *Pclo^wt/wt^*; *P* < 0.0001, Mann-Whitney-Wilcoxon test. This difference remains significant (*P* < 0.0001, Mann-Whitney-Wilcoxon test) when removing the two outliers from the control data set. The area of single-line and single spot-like Ca_V_1.3 clusters was also estimated by fitting a 2D-Gaussian function and was found to be smaller in *Pclo^gt/gt^* IHCs (*P* = 0.026 for single-line clusters, *P* = 0.028 for single spot-like clusters and *P* = 0.027 for all single structures; Mann-Whitney-Wilcoxon test; lines and spots pooled, represented in box plot in Fig 6E). This remained significant when removing the outlier from the control group (*P* = 0.03, Mann-Whitney-Wilcoxon test). The areas of ribbons and Ca_V_1.3 clusters showed a positive correlation for both *Pclo^wt/wt^ and Pclo^gt/gt^* (P_r_ = 0.53, *P* < 0.0001 for *Pclo^wt/wt^* and P_r_ = 0.44, *P* < 0.0001 for *Pclo^gt/gt^*, considering only synapses with single Ca_V_1.3 structures, Fig. 6E).

### Ca^2+^ current amplitude, voltage-dependence and kinetics remain unchanged upon piccolino disruption

We next investigated if piccolino disruption, in addition to altering the morphology of Ca^2+^ channel clusters, also affects Ca_V_1.3 Ca^2+^ channel physiology. We performed whole-cell perforated patch-clamp recordings of Ca^2+^ currents from IHCs of 2-4 weeks old rats in the presence of 5 mM [Ca]_e_ for enhanced signal to noise ratio. Step depolarizations of 20 ms from -87 to 68 mV in 5 mV step increments were applied to obtain current-voltage (IV) relations (Fig 7A). Ca^2+^ current amplitudes in *Pclo^gt/gt^* IHCs (-175.48 ± 12.87 pA, n_IHC_ = 20, N_animals_ = 13) appeared comparable to those in *Pclo*^wt/wt^ IHCs (-182.23 ± 10.21 pA, n_IHC_ = 20, N_animals_ = 13; *P* = 0.67, t-test, Fig 7A, Appendix Table 5). We did not find any statistically significant changes in the voltage-dependence of Ca^2+^ channel activation in piccolino deficient IHCs in contrast to what was observed in IHCs lacking synaptic ribbons (Jean et al., 2018) (Fig 7C). Activation and inactivation kinetics of Ca^2+^ channels also appeared unaltered (Fig 7B, D; see figure legend for description).

**Figure 7.**
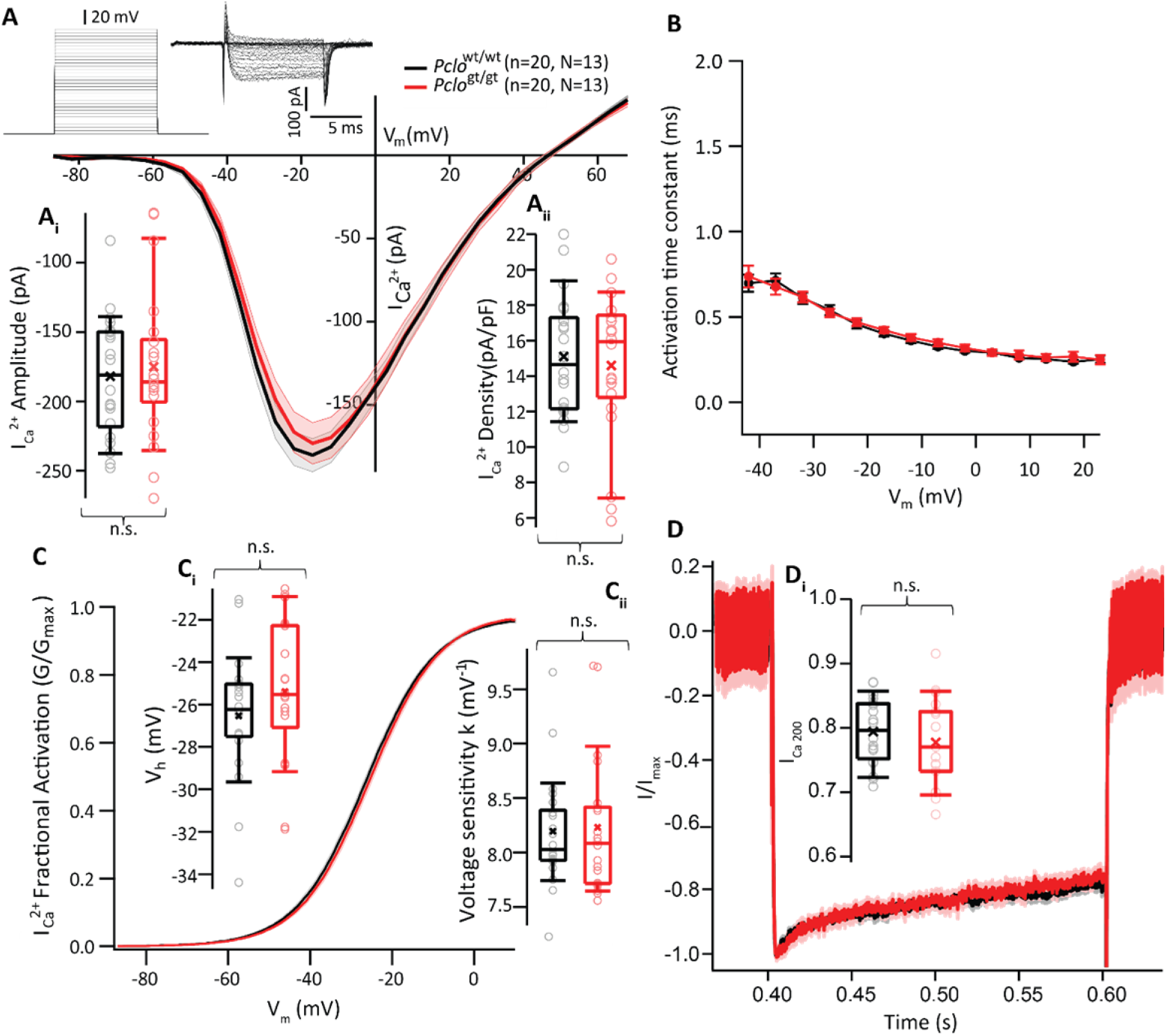
*Pclo^gt/gt^* IHCs show normal Ca^2+^ currents. **(A)** Current-voltage (IV) relations from whole-cell perforated patch clamp recordings of IHCs, [Ca]_e_ = 5 mM. Line represents mean current traces, shaded area represents ± SEM. **(A_i_)** A comparable whole-cell Ca^2+^ current amplitude (*P* = 0.67, t-test), and **(A_ii_)** current density (*P* = 0.68, t-test) was observed in IHCs of 2-4 weeks old *Pclo^wt/wt^* (n_IHC_ = 13, N_animals_ = 20) *Pclo^gt/gt^* (n_IHC_ = 13, N_animals_ = 20) rats. **(B)** A power exponential function was fitted on the first 5 ms of the current traces to obtain the activation time constant (mean ± SEM) of Ca^2+^ current at different voltages, which was unaltered. Estimation becomes less certain below -42 mV and above 23 mV, and hence these extremes were excluded from analysis. **(C)** A Boltzmann function was fitted to the current traces from **(A)** to derive the fractional activation of the whole-cell Ca^2+^ current. **(C_i_)** Voltage of half-maximal activation (V_h_) and **(C_ii_)** slope (k) of the Boltzmann fit do not show any statistically significant differences (*P* = 0.28 and *P* = 0.71 respectively, t-test). **(D)** Average peak amplitude-normalised Ca^2+^ currents in response to 200 ms depolarisations from -87 mV to -17 mV (shaded area represents ± SEM). Residual Ca^2+^ current **(D_i_)** show comparable inactivation in piccolino deficient IHCs; *P* = 0.22, Mann-Whitney-Wilcoxon test (n_IHC_ = 18, N_animals_ = 12 for *Pclo^wt/wt^* and n_IHC_ = 17, N_animals_ = 11 for *Pclo^gt/gt^*). Box plots show individual data points overlaid; mean values are shown as crosses.

**Appendix Table 5.**
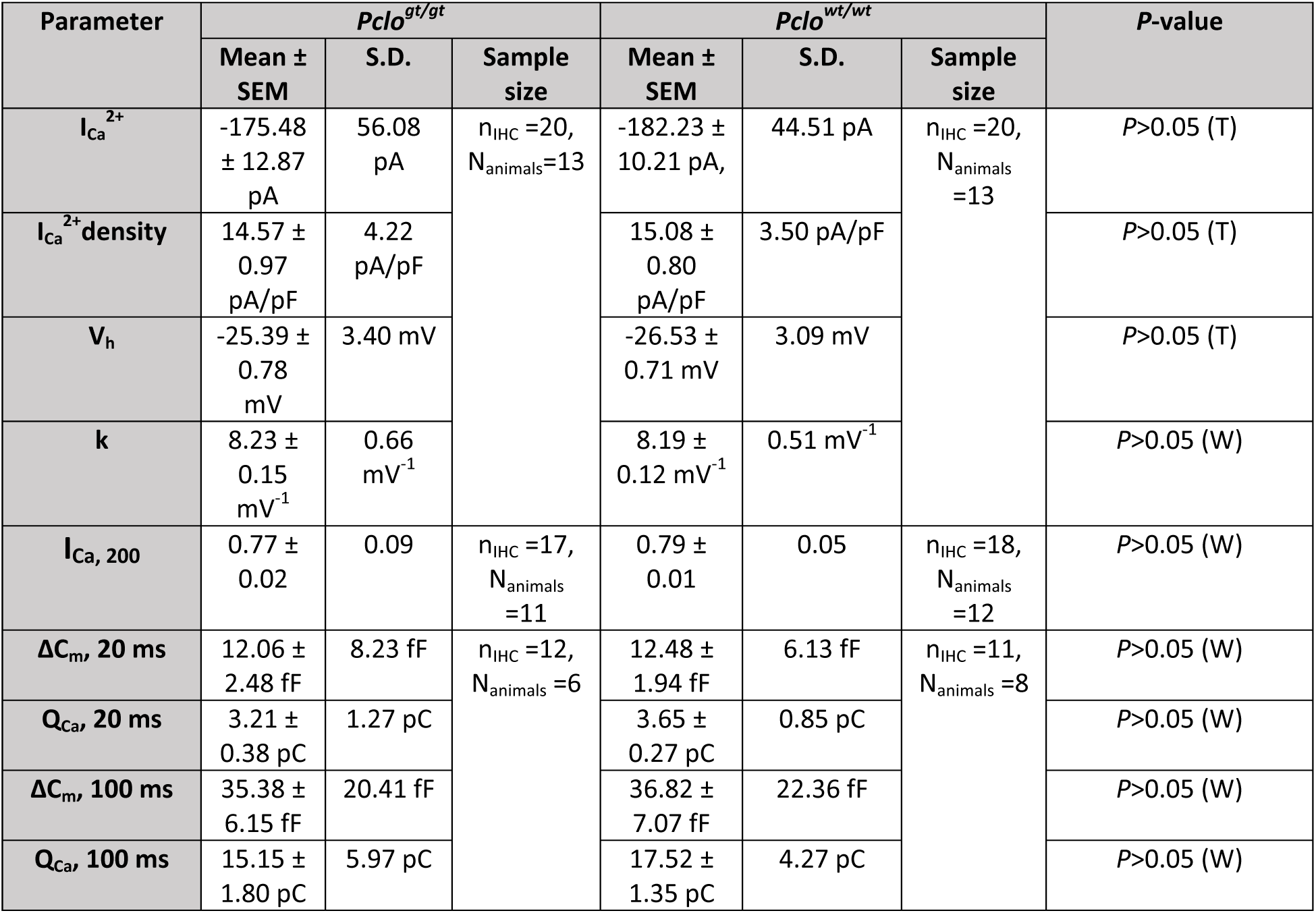
IHC physiology. Data is represented as mean ± SEM; n_IHC_ denotes number of IHCs, N_animals_ is number of rats used. Statistical analysis was performed using an unpaired two-sample *t* test (T), or using Mann-Whitney-Wilcoxon test (W). I_Ca_^2+^ = whole cell Ca^2+^ current, V_h_ = voltage of half maximal activation, k = voltage sensitivity of activation, I_Ca, 200_ = residual Ca^2+^ current at 200 ms depolarization, ΔC_m_ = change in exocytic membrane capacitance, Q_Ca_ = Ca^2+^ current integral.

### SV exocytosis and replenishment are unaltered in *Pclo^gt/gt^* IHCs

The synaptic ribbon plays a critical role in the exocytosis of the readily releasable pool of SVs (RRP) in retinal bipolar cells (Hull et al., 2006; Maxeiner et al., 2016; Snellman et al., 2011) and in rod photoreceptors (Grabner and Moser, 2021). The situation is more complex for cochlear IHCs lacking synaptic ribbons (ribbon-less): while the RRP was reduced in bassoon deficient IHCs (Buran et al., 2010; Khimich et al., 2005) it was normal in RIBEYE deficient IHCs (Becker et al., 2018; Jean et al., 2018). Some of this discrepancy can be attributed to additional effects of bassoon disruption e.g. on Ca^2+^ channel abundance and topography (Frank et al., 2010; Khimich et al., 2005; Neef et al., 2018) and compensatory synaptic transformations in the case of RIBEYE-deficiency (Jean et al., 2018). Here, we investigated if piccolino disruption impairs SV exocytosis and replenishment. We recorded exocytic membrane capacitance (ΔC_m_) in response to step depolarizations (to near the potential of maximal Ca^2+^ currents) using perforated patch clamp recordings in IHCs of 2-4 week old rats with 5 mM [Ca]_e_. ΔC_m_ in response to short and long depolarizations (recruiting the fast and sustained components of exocytosis respectively, Moser and Beutner, 2000) did not show any significant differences (Fig 8B, Appendix Table 5). To estimate the SV replenishment, we applied ten consecutive trains of 10 ms depolarization pulses (Fig 8D) and also performed paired pulse recordings (Fig 8E,F) as have been described before (Jean et al., 2018; Krinner et al., 2017). Again, we did not observe alterations in *Pclo^gt/gt^* IHCs which seems reminiscent of what was observed in ribbon-less IHCs of RIBEYE KO mice (Becker et al., 2018; Jean et al., 2018). Next, we analyzed the voltage-dependence of ΔC_m_ by stimulating IHCs with weak physiological depolarization pulses as RIBEYE KO IHCs showed a mild depolarized shift of exocytosis (Jean et al., 2018). In contrast, we do not observe any such alterations in exocytosis upon piccolino disruption (Fig 8G).

**Figure 8.**
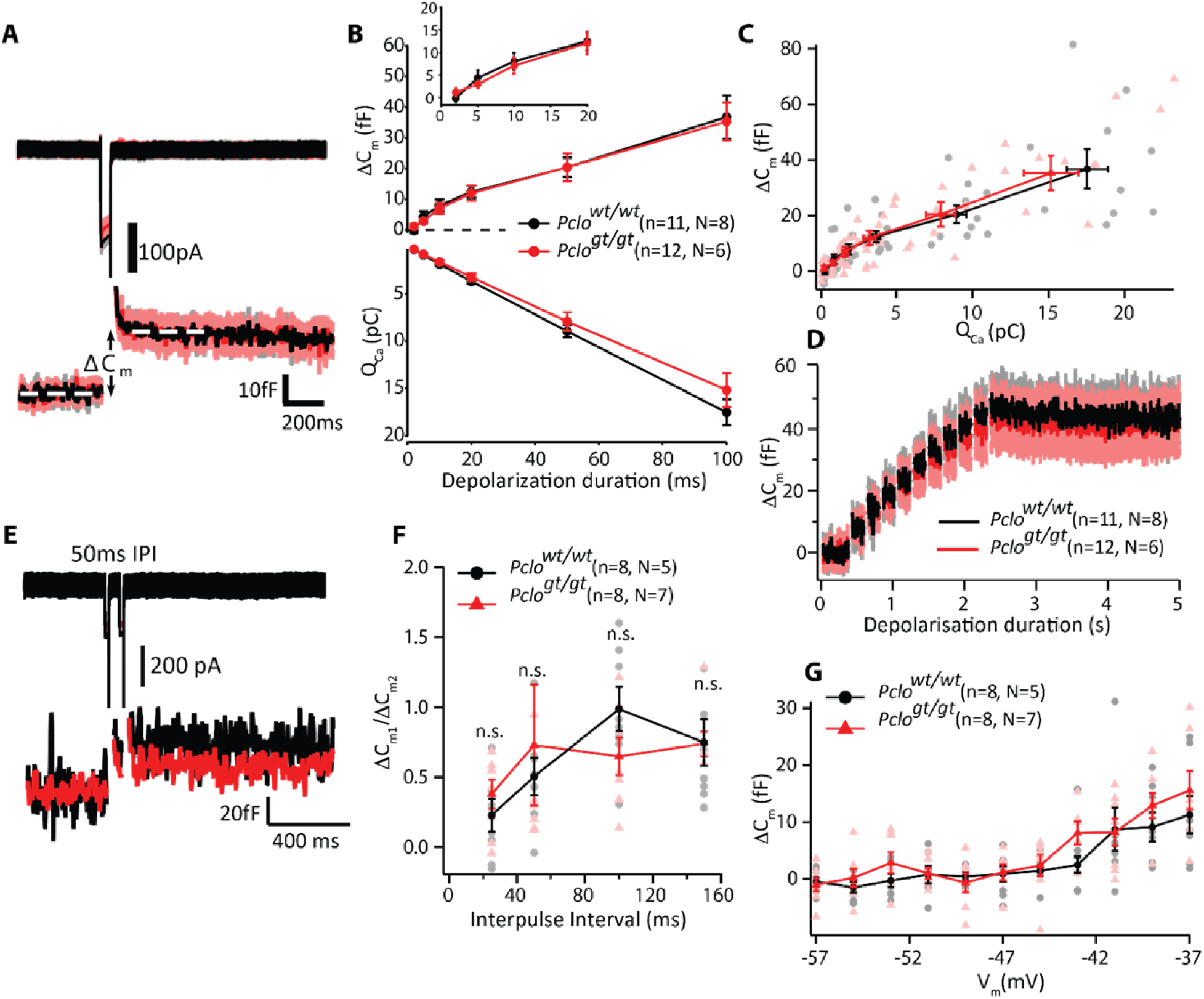
SV exocytosis and replenishment appear unaltered at the synapses of piccolino-deficient IHCs. **(A)** Average Ca^2+^ currents (upper panel) and corresponding changes in membrane capacitance (ΔC_m_, lower panel) in response to 50 ms depolarisations from -87 mV to -17 mV (shaded area represents ± SEM) from *Pclo^wt/wt^* IHCs (n_IHC_ = 11, N_animals_ = 8) and *Pclo^gt/gt^* IHCs (n_IHC_ = 12, N_animals_ = 6). **(B)** Mean ± SEM ΔC_m_ and the corresponding Ca^2+^ current integral (Q_Ca_) obtained in response to depolarisations of variable durations (2 to 100 ms) to -17 mV: normal phasic (≤ 20 ms) and sustained exocytosis in *Pclo^gt/gt^* IHCs (n_IHC_ = 12, N_animals_ = 6) in comparison to *Pclo^wt/wt^* IHCs (n_IHC_ = 11, N_animals_ = 8). Inset shows zoom-in for the first 20 ms. **(C)** ΔC_m_ vs. the corresponding Q_Ca_: normal apparent Ca^2+^ dependence of exocytosis (Mean ± SEM for each pulse duration are represented as darkened points, lightly shaded points represent individual IHCs). **(D)** Mean ΔC_m_ traces (shaded area represents ± SEM) in response to trains (10 stimuli) of 10 ms depolarisations from -87 to -17 mV: similar responses indicate intact SV replenishment for *Pclo^gt/gt^* IHCs (n_IHC_ = 12, N_animals_ = 6) and *Pclo^wt/wt^* (n_IHC_ = 11, N_animals_ = 8) IHCs. **(E)** Representative ΔC_m_ traces in response to pairs of 20 ms pulses separated by an inter-pulse interval (IPI) of 50 ms. **(F)** Paired-pulse ratios (ΔC_m2_/ΔC_m1_) at varying IPIs (25, 50, 100, and 150 ms) also indicate comparable rates of SV replenishment for *Pclo^gt/gt^* (n_IHC_ = 8, N_animals_ = 7) and *Pclo^wt/wt^* (n_IHC_ = 8, N_animals_ = 5) IHCs. **(G)** ΔC_m_ in response to 100 ms depolarisations of different potentials (Mean ± SEM and individual (lightly shaded points): no statistically significant difference, (n_IHC_ = 8, N_animals_ = 5 for *Pclo^wt/wt^*; n_IHC_ = 8, N_animals_ = 7 for *Pclo^gt/gt^*).

## Discussion

In the present study we analyzed the impact of genetic disruption of piccolino, the short isoform of the multidomain protein piccolo/aczonin on the structure and function of IHC ribbon synapses. Using a multidisciplinary approach, we identified a mild synaptic hearing impairment and roles of piccolino for ribbon morphology, SV complement and Ca^2+^ organization at the IHC AZ. Intriguingly the observed subtle structural alterations seemed to affect only a subset of AZs that, however, distributed throughout the synaptic IHC pole regardless of position along the modiolar-pillar axis. Moreover, we failed to detect alterations of presynaptic function at least on the whole IHC level which contrasts with a mild sound encoding deficit observed on the systems level. Together our results indicate that piccolino is required for normal structure and function of IHC synapses.

### Mild functional and structural impairment upon piccolino disruption

Genetic disruption of piccolino in rats (10-weeks-old *Pclo^gt/gt^*) resulted in a mild hearing deficit with elevated ABR thresholds for middle and high frequency tone bursts and a significant reduction of the amplitude of the first ABR wave reflecting an impairment in synchronous firing of SGNs. Previous studies in piccolo KO mice (Mukherjee et al., 2010) reported normal sound thresholds and a reduced wave II amplitude (Butola et al., 2017). In these mutants, the piccolino splice variant was not affected (Butola et al., 2017), thus normal sound encoding on the level of ribbon synapses was expected, given that piccolino is the only isoform present at ribbon synapses (Regus-Leidig et al., 2013). Recently, Li and co-workers reported findings in mice with complete loss of piccolo and piccolino demonstrating functional impairments of the retina while hearing seemed unaffected with normal ARBs up to 8 months of age (Li et al., 2021). Our findings on piccolino gene-trap rats contrast the normal hearing in KO mice for which we currently do not have an explanation.

Moreover, using STED and electron microscopy, we observed several alterations of ribbon synapse morphology in cochlear IHCs of *Pclo^gt/gt^* rats. A fraction of the Ca_V_1.3 Ca^2+^ channel clusters appeared fragmented into smaller patches. Furthermore, focusing on single spot-like and single line-shaped Ca_V_1.3 Ca^2+^ channel clusters we found them to be smaller, as were synaptic ribbons, while PSDs were enlarged. Electron microscopy uncovered two morphologically distinct AZ categories at mutant synapses: category 1 closely resembled control AZ, while ribbons of category 2 AZs were strikingly smaller and completely lacked SVs at the membrane-distal part of the ribbon. To our surprise, these structural alterations were not accompanied by functional changes on the whole IHC level: Ca^2+^ currents and exocytosis of *Pclo^gt/gt^* IHCs were comparable to the controls. It will be interesting in future studies to study synaptic function at the level of single synapses in order to test the hypothesis that postulated functional deficits of category 2 AZs were masked by category 1 AZs. Indeed, category 1 AZs were found to be even larger compared to littermate controls with an elevated number of SVs. Together, the presence of both, enlarged category 1 AZs and the small category 2 AZs, yielded normal Ca^2+^ currents and exocytic function at the whole IHC level. Further, the enlarged PSDs, which we observed in the *Pclo^gt/gt^* rats might contribute to compensate for presynaptic structural deficits. Such enlarged PSDs were previously observed in piccolo and bassoon mutants in conventional synapses (Mukherjee et al., 2010). *Ex vivo* patch-clamp recordings from IHCs of higher frequency places of the cochlea as well as *in vivo* recordings of single SGNs in future studies might help to further elucidate the relationship of exocytosis, which could differ in its dependence on piccolino along the tonotopic axis, and neural sound encoding.

### A role of piccolino as a structural determinant for ribbon synapses

Previously, it was shown for several species that morphological and functional attributes of IHC ribbon synapses display a spatial gradient along the modiolar (or neural) – pillar (or abneural) axis. IHC AZs vary in ribbon size (Kantardzhieva et al., 2013; Liberman, 1980; Meyer et al., 2009; Michanski et al., 2019; Ohn et al., 2016; Payne et al., 2021), number, voltage-dependence of activation and coupling of SVs to Ca^2+^ channels (Frank et al., 2009; Ohn et al., 2016; Özçete and Moser, 2021). Such synaptic diversity offers an exciting candidate mechanism for how IHCs might decompose the full range of sound intensity information into complementary neural codes in SGNs that have long been known to differ in spontaneous and sound-evoked firing (Moser et al., 2019; Rutherford and Moser, 2016). Classically SGNs have been categorized into three functional subtypes: low, medium, and high spontaneous rate (SR) SGNs (Kiang et al., 1965; Liberman, 1978; Sachs and Abbas, 1974; Taberner and Liberman, 2005; Winter et al., 1990). Recently, analysis of molecular SGN profiles has led to the identification of three molecular SGN subtypes type 1a-c which were suggested to relate to the functional SGN subtypes (Li et al., 2020; Petitpré et al., 2018; Shrestha et al., 2018; Sun et al., 2018) and might also instruct properties of the corresponding presynaptic IHC AZ (Sherrill et al., 2019). Last but not least, differences in efferent innervation of the peripheral SGNs neurites (e.g. Hua et al., 2021; Liberman et al., 1990) might contribute to the synaptic and neurophysiological differences of SGNs (Ruel et al., 2001; Yin et al., 2014). How the various mechanisms interplay to the collective sound encoding of SGNs has yet to be elucidated. Specifically, how IHCs diversify their AZs remains an exciting research question. Specifically, identifying molecular pathways and determinants setting the specific structure and function is a key task.

The present finding of two categories of presynaptic AZs in *Pclo^gt/gt^* IHCs despite the given overall AZ variance provides an interesting insight into how individual AZ proteins can determine structure. A structural role of piccolino at ribbon synapses was previously demonstrated in the retina (Regus-Leidig et al., 2014). It is conceivable that its interaction with RIBEYE plays an important role in maintaining synapse architecture (Müller et al., 2019). Already the reduction of piccolino resulted in structural changes of plate-like photoreceptor ribbons to smaller, more oval shaped ones (Regus-Leidig et al., 2014). Müller et al. discussed the option that RIBEYE together with piccolino organizes the plate-like structure of the synaptic ribbon in the retina, specifically for rod photoreceptors and rod bipolar cells. Since IHC ribbon synapses are more oval in their shape, the authors suggested that probably a different RIBEYE-piccolino ratio might exist at IHC ribbon synapses, which would result in their oval structure (Müller et al., 2019). Grading piccolino abundance across the AZs of individual IHCs might likewise contribute to different morphologies of their ribbons. However, whether different ratios of piccolino/RIBEYE prevail at the AZs of individual IHCs remains to be investigated in future experiments. The present findings on piccolino deficient IHCs did not point to preferred position along the modiolar-pillar axis of the AZ categories, which would not seem to support a major role of piccolino in shaping synaptic heterogeneity in IHCs. We speculate that the ribbon of category 2 AZ found upon disrupting piccolino might represent a sort of basic IHC ribbon version, which is then modulated by the amount of piccolino, while category 1 might reflect a compensatory potential specific to a subset of synapses. Addressing functional differences between the AZ categories in *Pclo^gt/g^* IHCs will require future single synapse analysis (Goutman and Glowatzki, 2007; Özçete and Moser, 2021).

### Normal synaptic vesicle tethering at piccolino-deficient AZs

Piccolino was shown to bind to RIBEYE, while its N-terminus faces the cytoplasm and was suggested to regulate steps in the SV cycle by binding to other presynaptic proteins (Müller et al., 2019). In line with previous studies for photoreceptor ribbon synapses (Limbach et al., 2011; Müller et al., 2019), our immunogold labelings of rodent ribbon synapses confirmed that piccolino localizes to synaptic ribbons of IHCs (the present study, (Michanski et al., 2019)). Moreover, ribbons of *Pclo^gt/gt^*category 2 AZs were void of membrane distal SVs, which could point either to a role of piccolino in SV tethering or in enabling a tethering-relevant ribbon part. Importantly, normal ribbon-tether number of those remaining RA-SVs at category 2 AZs argue against an essential role of piccolino in tethering SVs to the ribbon. Nonetheless, the increased distance of SVs to the ribbon surface of AZs could indicate a contribution of piccolino in attracting SVs. Moreover, a previous study has shown that a 110 kDa piccolino fragment is still detectable in *Pclo^gt/gt^* rat retina (Müller et al., 2019), which could exert some residual function. For example, the interaction site for the actin/dynamin-binding protein Abp1 is likely still present in the *Pclo^gt/gt^* mutant rats, while the RIBEYE interaction site, which is located in the C-term region of piccolino, is eliminated (Müller et al., 2019). Fenster et al. showed the interaction between piccolo and Abp1, which also binds to both F-actin and the GTPase dynamin. This led to the hypothesis that piccolo localizes Abp1 to AZs to create a functional connection between the dynamic actin cytoskeleton and SV recycling at conventional synapses (Fenster et al., 2003), the relevance of which for ribbon synapses remains to be tested.

### Key molecular organizers of the ribbon-type AZs in IHCs

Bassoon and piccolo are important scaffold proteins at AZs with partially overlapping functions (review in Gundelfinger et al., 2015). Disruption of piccolo resulted in normal sized AZs but fewer SVs in AZs of endbulb of Held synapses (Butola et al., 2017). In AZs of rat IHCs, the disruption of piccolino had several structural consequences (Fig 9). Mutant AZs showed only single ribbons and generally fewer MP-SVs. Moreover, a subset of AZs with smaller ribbons was found, which lack SVs at their membrane distal side. The Ca^2+^ channel clusters appeared fragmented, resembling what has been shown for IHC AZs in RIBEYE mutants (Jean et al., 2018). Altered localization and reduced number of Ca^2+^ channels were previously reported for RIM2 (Jung et al., 2015a) and RIM-BP2 (Krinner et al., 2017) deficient IHC AZs. RIM2 and RIM-BP2 positively regulate the number of synaptic Ca_V_1.3 channels at IHC AZs and promote fast SV recruitment to the RRP. Loss of the ribbon due to disruption of RIBEYE resulted in subtle functional impairments in IHCs and hearing (Fig 9) (Becker et al., 2018; Jean et al., 2018), while in rod photoreceptors RIBEYE disruption dwarfed the RRP to a third (Grabner and Moser, 2021). Maintained presynaptic IHC function in ribbonless IHCs was attributed to partial compensation that encompasses several small bassoon-positive presynaptic densities at ribbon-less AZs tethering a large number of SVs (Jean et al., 2018). This raised the question if ribbons are dispensable for the function of IHC AZs as long as bassoon remains present. Piccolino lacks the interactions site to bassoon (Regus-Leidig et al., 2013), and different from conventional synapses, piccolino and bassoon segregate at ribbon-type AZs (Dick et al., 2001; Khimich et al., 2005; Limbach et al., 2011; Michanski et al., 2019; tom Dieck et al., 1998; Wong et al., 2014). In contrast to the mild sensory coding phenotypes of piccolino mutants (the present study, (Li et al., 2021; Müller et al., 2019)), disruption of bassoon strongly alters transmission at afferent synapses of retina and cochlea (Fig 9). The deletion of exon 4 and 5 (*Bsn^ΔEx4/5^*) resulted in detached synaptic ribbons in photoreceptors and IHCs and impaired sensory coding assayed at the systems and synaptic levels (Dick et al., 2003; Khimich et al., 2005). In addition, to the reduced RRP, *Bsn^ΔEx4/5^* IHC AZs showed fewer and mislocalized Ca^2+^ channels (Frank et al., 2010; Khimich et al., 2005; Neef et al., 2018).

**Figure 9:**
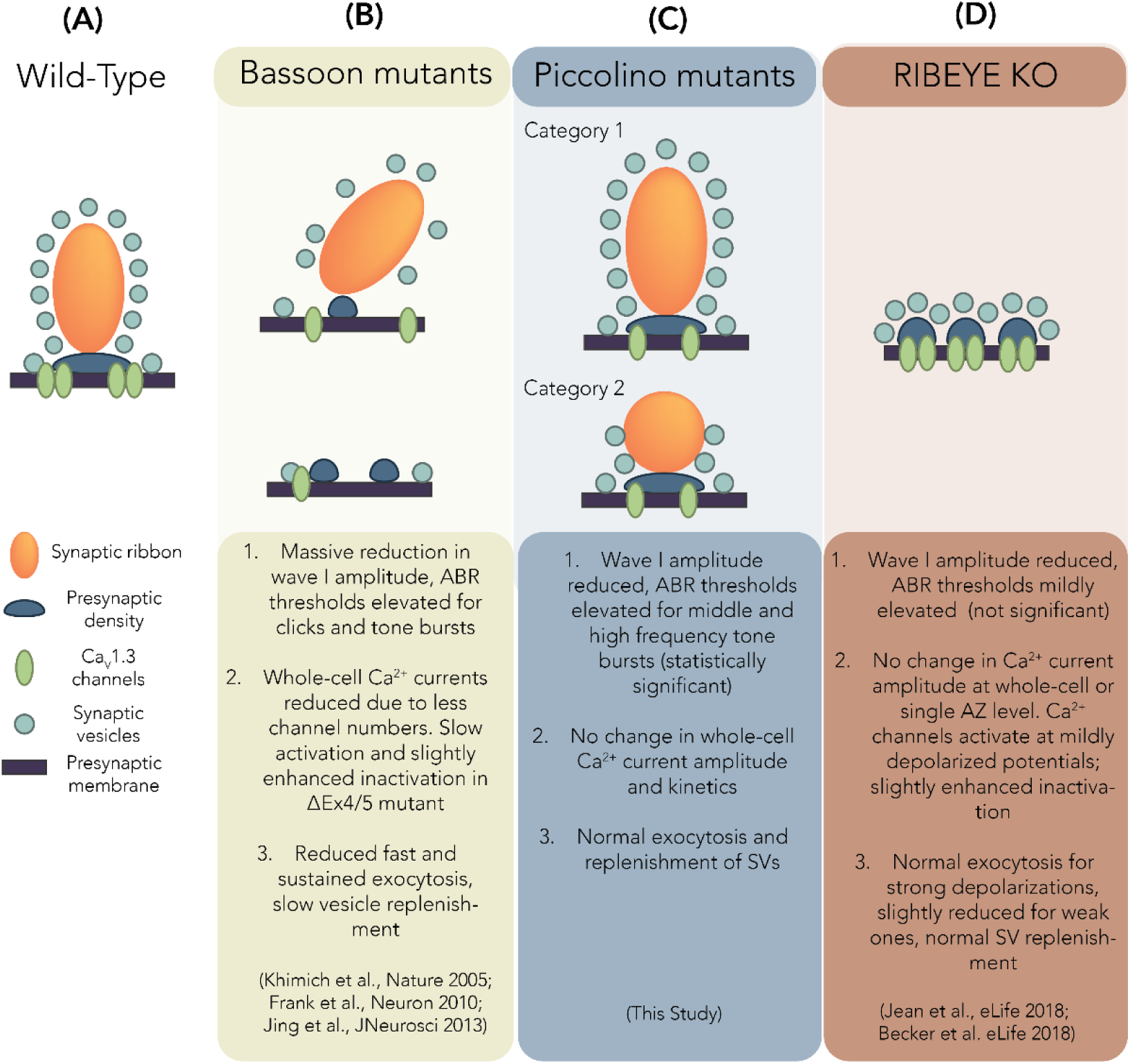
Summary of key structural and functional observations at IHC AZs upon genetic perturbation of bassoon, piccolino and RIBEYE. (A) Wild-type IHC synaptic ribbons drawn for reference; these are typically ellipsoid shaped and anchored to the presynaptic membrane via the presynaptic density. **(B)** Disruption of bassoon causes ribbon anchorage defects, resulting in ribbon-less AZs (lower panel, representing partial deletion ΔEx4/5 mutants) or loosely anchored ribbons (top panel, representing the predominant form in bassoon gene-trap mutants). **(C)** Disruption of piccolino results in two categories of AZs: category 1 wild-type-like ribbons and roundish, small ribbons for category 2 AZs that lack SVs at their membrane-distal apex. **(D)** RIBEYE deficient IHCs show ribbon-less AZs with multiple smaller presynaptic densities and SV assemblies feeding into the same postsynapse. Ca^2+^ channel clusters appear smaller and fragmented in IHCs of bassoon, piccolino and RIBEYE mutants.

It has remained challenging to disentangle effects of ribbon-loss and overall AZ alterations upon bassoon disruption (Frank et al., 2010; Jing et al., 2013), in particular given the mild functional deficits in ribbon-less IHCs of RIBEYE KOs (Becker et al., 2018; Jean et al., 2018). We suggest that bassoon - as the major component of the presynaptic density (Wong et al., 2014) - is the main organizer at IHC ribbon synapse AZs and might even be able to tether SVs to the presynaptic density. Piccolino and RIBEYE likely jointly determine ribbon shape and size and might thus finetune exocytosis at IHC ribbon synapses.

## Material and Methods

### Animals

We used a piccolo gene trap (*Pclo^gt/gt^*) rat strain, which is described in detail in (Medrano et al., 2020). Briefly, transposon mutagenesis resulted in the integration of a transposon element into exon 3 of the *PCLO* gene, leading to a stop in the reading frame. *Pclo^gt/gt^* rats and wild-type littermate controls (*Pclo^wt/wt^*), from heterozygous breeding of either sex between 2 weeks and 11 months, were deeply anesthetized with CO_2_ and sacrificed by decapitation for immediate dissection of the cochleae. All experiments complied with national animal care guidelines and were approved by the University of Göttingen Board for Animal Welfare and the Animal Welfare Office of the State of Lower Saxony.

### Systems physiology

ABR and DPOAE were performed for 2-months old rats as described before (Jing et al., 2013; Strenzke et al., 2016). Animals were anesthetized intraperitoneally with a combination of ketamine (125 mg/kg) and xylazine (2.5 mg/kg), and the heart rate was constantly monitored to control the depth of anesthesia. The core temperature was maintained constant at 37°C using a rectal temperature-controlled heat blanket (Hugo Sachs Elektronik–Harvard Apparatus). For stimulus generation, presentation, and data acquisition, we used the TDT III Systems (Tucker Davis Technologies) run by BioSig32 software (Tucker Davis Technologies). Sound pressure levels (SPL) are provided in decibels SPL root mean square (RMS) (tonal stimuli) or decibels SPL peak equivalent (clicks) and were calibrated using a ¼ inch Brüel and Kjær microphone (model 4939). Tone bursts (4/8/12/16/24/32 kHz, 10 ms plateau, 1 ms cos^2^ rise/fall) or clicks of 0.03 ms were presented at 20 Hz in the free field ipsilaterally using a JBL 2402 speaker. The difference potential between vertex and mastoid subdermal needles was amplified (50,000 times), filtered (low pass, 4 kHz; high pass, 100 Hz) and sampled at a rate of 50 kHz for 20 ms, 2 × 2000 times to obtain two mean ABRs for each sound intensity. Hearing threshold was determined with 10 dB precision as the lowest stimulus intensity that evoked a reproducible response waveform in both traces by visual inspection. For DPOAEs, a 24-bit sound card and the MF1 speaker system (Tucker David Technologies) were used to generate two primary tones (f2/f1 ratio: 1.2). Primary tones were coupled into the ear canal by a custom-made probe containing an MKE-2 microphone (Sennheiser) and adjusted to an intensity of 60 dB sound pressure level at the position of the ear drum as mimicked in a mouse ear coupler. The microphone signal was amplified (DMX6Fire; Terratec) and analyzed by fast Fourier transformation using custom software (Matlab, mathworks).

### Immunohistochemistry and Imaging

Cochleae from 2-3 weeks old (postnatal day P14 - P27) and 2 months old rats were dissected in ice-cold HEPES Hank’s solution containing (in mM): 5.26 KCl, 141.7 NaCl, 0.5 MgSO_4_-7H_2_O, 10 HEPES, 1 MgCl_2_, 11.1 D-glucose and 3.42 L-glutamine, pH adjusted to around 7.2, osmolality ∼300 mOsm/kg. Fixation was performed by perfusing the cochleae with 4% formaldehyde (in PBS) for 60 min on ice, while for staining Ca^2+^ channel clusters, a shorter fixation of 5-10 min was performed. For cochleae from 2 months old rats, decalcification was additionally performed using Morse’s solution (10% sodium citrate, 22.5% formic acid) for 15-20 min. The organ of Corti was dissected and washed three times in PBS at room temperature. Blocking and permeabilisation of the tissue was performed with GSDB (goat serum dilution buffer: 16% normal goat serum, 450 mM NaCl, 0.3% Triton X100, 20 mM phosphate buffer, pH ∼7.4) for 1 h at room temperature. Samples were then incubated with primary antibodies (diluted in GSDB, refer to table 1) overnight at 4°C and were washed three times for 10 min in wash buffer (450 mM NaCl, 0.3% Triton X 100, 20 mM phosphate buffer, pH ∼ 7.4). This was followed by incubation with respective secondary antibodies (diluted in GSDB, refer to table 1) for 1 h in a light-protected wet chamber. Finally, the samples were washed three times for 10 min in wash buffer and in 5 mM phosphate buffer for 5 min, before mounting onto glass slides with a drop of fluorescence mounting medium (*Mowiol* 4-88, Carl Roth) and covered with thin glass coverslips. Images were acquired in confocal/STED mode using an Olympus IX83 inverted microscope combined with an Abberior Instruments Expert Line STED microscope (Abberior Instruments GmbH). We employed lasers at 488, 561, and 633 nm for excitation and a 775 nm (1.2W) laser for stimulated emission depletion. 1.4 NA 100X or 20X oil immersion objectives were used. Confocal stacks were acquired using *Imspector* Software (pixel size = 80 X 80 nm along xy, 200 nm along z). For 2D-STED images, a pixel size of 15 X 15 nm (in xy) was used. The acquired images were z-projected with NIH ImageJ software and adjusted for brightness and contrast. Organs of Corti from both *Pclo^wt/wt^* and *Pclo^gt/gt^* were always processed in parallel using identical staining protocols, laser excitation powers and microscope settings.

**Table 1:** List of antibodies:

| Antibody | Host specie | Company | Dilution | Identifier |
| --- | --- | --- | --- | --- |
| Anti-parvalbumin | Guinea pig (polyclonal) | Synaptic Systems | 1:200 | 195 004 |
| Anti-CtBP2 | Mouse (monoclonal IgG1) | BD Biosciences | 1:200 | 612044 |
| Anti-piccolo | Rabbit (polyclonal) | Synaptic Systems | 1:200 | 142 113 |
| Anti-bassoon | Chicken (polyclonal) | Synaptic Systems | 1:200 | 141 016 |
| Anti-homer1 | Rabbit (polyclonal) | Synaptic Systems | 1:200 | 160 002 |
| Anti-Ca<sub>v</sub>1.3 (CACNA1D) | Rabbit (polyclonal) | Alomone Labs | 1:100 | ACC 005 |
| Alexa Fluor 488 conjugated anti-guinea pig | Goat (polyclonal) | Invitrogen | 1:200 | A11073 |
| Alexa Fluor 568 conjugated anti-rabbit | Goat (polyclonal) | Thermo Fisher | 1:200 | A11011 |
| STAR635p conjugated anti-mouse | Goat (polyclonal) | Abberior | 1:200 | 2-0002-007-5 |
| Alexa Fluor 488 conjugated anti-rabbit | Goat (polyclonal) | Invitrogen (MoBiTec) | 1:200 | A11008 |
| Alexa Fluor 568 conjugated anti-chicken | Goat (polyclonal) | Abcam | 1:200 | Ab175711 |
| Alexa Fluor 488 conjugated anti-chicken | Goat (polyclonal) | Invitrogen | 1:200 | A11039 |
| STAR580 conjugated anti- | Goat (polyclonal) | Abberior | 1:200 | 2-0002-005-1 |
| mouse |  |  |  |  |
| STAR635p conjugated anti-rabbit | Goat (polyclonal) | Abberior | 1:200 | 2-0012-007-2 |

### Focused ion beam-scanning electron microscopy (FIB-SEM)

Enhanced en bloc staining for FIB-SEM samples of 2 *Pclo^wt/wt^* and 2 *Pclo^gt/gt^*rats was performed according to Deerinck et al. (Deerinck et al., 2018) and to our previous studies (Jean et al., 2020; Michanski et al., 2019). Organs of Corti, within the apical turn of the cochlea from 2-3 months old rats, were isolated in ice-cold HEPES Hank’s solution (5.36 mM KCl (746436, Sigma-Aldrich, Germany), 141.7 mM NaCl (746398, Sigma-Aldrich, Germany), 10 mM HEPES (H3375, 006K5424, Sigma-Aldrich, Germany), 0.5 mM MgSO_4_-7H_2_O (Sigma-Aldrich, Germany), 1 mM MgCl_2_ (M2670, Sigma-Aldrich, Germany), 2 mg/ml D-glucose (G8270-1KG, Sigma-Aldrich, Germany), 0.5 mg/ml L-glutamine (G3126-100G, #SLBS8600, Sigma-Aldrich, Germany) and was adjusted to pH 7.2, ∼300 mmol/kg). After the dissection, organs were immediately fixed with 4% paraformaldehyde (0335.1, Carl Roth, Germany) and 0.5% glutaraldehyde (G7651, Sigma-Aldrich, Germany) in PBS (P4417, Sigma-Aldrich, Germany; pH 7.4) for 1 h on ice followed by a second fixation step overnight with 2% glutaraldehyde in 0.1 M sodium cacodylate buffer (v/v, pH 7.2) at 4°C. The next day, fixed tissues were treated with a 1.5% potassium ferrocyanide (25154-10, EMS) and 4% osmium tetroxide solution (75632.5ml, Sigma-Aldrich, Germany; v/v in 0.1 M sodium cacodylate buffer) for 1 h on ice. Specimens were then briefly washed 5 times in distilled water and placed in a thiocarbohydrazide (w/v in distilled water) solution for 20 min followed by additional 5 brief washing steps in distilled water at room temperature (RT). Next, a second incubation into 2% osmium tetroxide (v/v in 0.1 M sodium cacodylate buffer) for 2 h at RT followed with subsequent 5 brief washing steps in distilled water before the samples were placed in 2.5% uranyl acetate (v/v in distilled water) overnight at RT and in darkness. The following day, samples were briefly washed 5 times in distilled water at RT and contrasted with Reynold’s lead citrate (Reynolds, 1963) for 30 min at 60°C to be subsequently washed once again in distilled water, dehydrated in increasing ethanol concentrations (30%, 50%, 70%, 95% and 100%), infiltrated and finally embedded in Durcupan (25%, 50%, 75% Durcupan in Acetone for 1 h each and 100% Durcupan overnight at RT and for another 2 h with fresh 100% Durcupan on the following day; 44610, Sigma-Aldrich, Germany) to get polymerized for at least 48 h at 60°C. The cured blocks were then trimmed with a 90° diamond trimming knife (Diatome AG, Biel, Switzerland), attached to SEM stubs (Science Services GmbH, Pin 12.7 mm x 3.1 mm) using a silver filled epoxy (Epoxy Conductive Adhesive, EPO-TEK EE 129-4; EMS) and polymerized at 60° overnight. Afterwards, samples on SEM stubs were coated with a 10 or 15 nm platinum or gold layer using the sputter coating machine EM ACE600 (Leica Microsystems) at 30 mA current to be finally placed into the Crossbeam 540 FIB-SEM (Carl Zeiss Microscopy GmbH) and positioned at 54°. A 400 or 500 nm carbon or platinum layer was deposited on top of the regions of interest and the Atlas 3D (Atlas 5.1, Fibics, Canada) software was used to collect the acquired image dataset. Specimens were exposed to the ion beam driven with a 30 nA current while a 7 or 15 nA current was applied to polish the surface. Images were acquired at 1.5 kV (1000 pA) using the ESB detector (450 V ESB grid, pixel size x/y 3 or 5 nm) in a continuous mill and acquire mode using 700 pA or 1.5 nA for the milling aperture (z-step 5 or 10 nm). For subsequent post processing, data were aligned using the Plugin “Linear Stack Alignment with SIFT”, inverted and cropped in Fiji. Depending on the dataset properties, a Gaussian blur, local contrast enhancement using a CLAHE plugin in Fiji, and a binning by 2 in x/y was applied (Schindelin et al., 2012).

### High-pressure freezing (HPF) and Freeze-substitution (FS)

High-pressure freezing was essentially performed as described previously (Chapochnikov et al., 2014; Jung et al., 2015a). In brief, the apical turn cochlear organs from 1-2 months old *Pclo^wt/wt^* and *Pclo^gt/gt^* rats were dissected in ice-cold HEPES Hank’s solution and mounted onto type A specimen carriers (Leica Microsystems, Wetzlar, Germany; 3 mm in diameter and 0.2 mm in depth). The 1-hexadecene (Sigma-Aldrich, Germany) coated flat side of the type B carriers (Leica Microsystems, Wetzlar, Germany; 3 mm in diameter and 0.1 mm in depth) was then placed onto the sample containing type A carrier. The assembled carriers were loaded into the middle plates of the high-pressure freezing sample holder and excess liquid was removed with filter paper. Afterwards, the sample holder was assembled and loaded into the HPM100 (Leica Microsystems, Wetzlar, Germany) to cryofix the tissues and store them in liquid nitrogen until further processing. Subsequently, the high-pressure frozen samples were freeze-substituted using the EM AFS2 (Leica Microsystems, Wetzlar, Germany) machine. In brief, organs were incubated in 0.1% (w/v) tannic acid in acetone at -90°C for 4 days followed by 3 washing steps in acetone at -90°C for 1 h, respectively. Next, 2% (w/v) osmium tetroxide in acetone was applied and incubated at -90°C for 7 h. During the following 33.4 h the temperature gradually rose up to 4°C and osmium tetroxide in acetone was removed, samples were washed in acetone 3 times (1 h each) and brought to RT. Finally, samples were infiltrated and embedded in epoxy resin (Agar-100 kit, Plano, Germany; epoxy/acetone 1:1 3-6 h; 100% epoxy resin overnight and 3-6 h on the next day) to get polymerized for at least 48 h at 70°C.

### Conventional embedding

Cochlea organs from 3 months old *Pclo^wt/wt^* and *Pclo^gt/gt^* rats (one animal each) were dissected as described above. Subsequently, organs were fixed immediately after dissection with 4% paraformaldehyde (0335.1, Carl Roth, Germany) and 0.5% glutaraldehyde (G7651, Sigma, Germany) in PBS (P4417, Sigma, Germany; pH 7.4) for 1 h on ice followed by a second fixation step overnight with 2% glutaraldehyde in 0.1 M sodium cacodylate buffer (v/v, pH 7.2) at 4°C. Next, specimens were washed in 0.1 M sodium cacodylate buffer and treated with 1% osmium tetroxide (75632.5ml, Sigma, Germany; v/v in 0.1 M sodium cacodylate buffer) for 1 h on ice followed by further sodium cacodylate buffer and distilled water washing steps. After the *en bloc* staining with 1% uranyl acetate (8473, Merck, Germany; v/v in distilled water) for 1 h on ice, samples were briefly washed in distilled water, dehydrated in an ascending concentration series of ethanol (30%, 50%, 70%, 95% and 100%), infiltrated and embedded in epoxy resin (R1140, AGAR-100, Plano, Germany) to get finally polymerized for at least 48 h at 70°C.

### Immunogold pre-embedding

The Triton X immunogold pre-embedding protocol was applied according to our previous study (Michanski et al., 2019). In brief, cochlea organs from 10-11 months old *Pclo^wt/wt^* and *Pclo^wt/gt^* rats (N_animals_ = 2 for each genotype) were dissected as mentioned above and fixed with 2% paraformaldehyde and 0.06% glutaraldehyde in PEM (0.1 M PIPES: P1851-500g, Sigma, Germany; 2 mM EGTA: E3889, Sigma, Germany; 1 mM MgSO_4_ x 7 H_2_O, v/v) for 90 min on ice. Afterwards, samples were washed in PEM and blocked for 1 h in 2% bovine serum albumin (BSA; 900099, Aurion)/ 3% normal horse serum (NHS; VEC-S-200, Biozol, Germany) in 0.02% PBST (0.02% Triton X-100 (X100-500ml, Sigma, Germany) diluted in PBS, v/v). Next, samples were incubated with the anti-rabbit piccolo primary antibody (polyclonal, Synaptic Systems: 142113; 1:200 diluted in 0.02% PBST), detecting the long and the short (piccolino) isoform of piccolo for 1 h at RT and overnight at 4°C. Subsequently, specimens were washed with 0.02% PBST and incubated for 2 h with the 1.4 nm gold-coupled anti-rabbit secondary antibody (Nanogold-anti-rabbit, Nanoprobes; 1:30 diluted in 0.02% PBST) followed by another washing step in 0.02% PBST for 30 min and overnight at 4°C. Next day, after further washing steps in 0.02% PBST, samples were post-fixed with 2% glutaraldehyde in PBS (v/v) for 30 min and briefly washed in distilled water. For silver enhancement, the HQ Silver-enhancement kit (Nanoprobes) was used for 3 min in the dark and specimens were briefly washed four times. Further fixation was obtained by the treatment with 2% osmium tetroxide (v/v in 0.1 M cacodylate buffer) for 30 min followed by one washing step in distilled water for 1 h and two washing steps in distilled water for 30 min, respectively. Finally, samples were dehydrated in an ascending concentration series of ethanol (30%, 50%, 70%, 95% and 100%), infiltrated and embedded in epoxy resin (R1140, AGAR-100, Plano, Germany) and polymerized for at least 48 h at 70°C.

### Ultrathin-sectioning and post-staining

The polymerized blocks were trimmed into a pyramidal shape to remove excess resin and 70 nm ultrathin sections were cut with a 35° diamond knife (Diatome AG, Biel, Switzerland) using an EM UC7 (Leica Microsystems, Wetzlar, Germany) ultramicrotome in order to check for the correct region as well as the structural preservation. Ultrathin sections were collected on 1% formvar-coated copper slot grids (Athene, 3.05 mm diameter, 1 x 2 mm; G2500C, Plano). For electron tomography, 250 nm semi-thin sections were obtained and collected on 1% formvar-coated mesh grids (100 mesh; Athene, 3.05 mm diameter; G2410C, Plano). For both sectioning techniques, post-staining was performed using UranyLess (22409, EMS, Hatfield, PA) for 20 min followed by several brief washing steps with distilled water.

### Transmission electron microscopy and electron tomography

For conventional embedded samples, immunogold pre-embedded samples and to first check for the quality of the cryo-fixed tissues and the region of interest, 2D electron micrographs were taken from 70 nm ultrathin sections at 80 kV using a JEM1011 transmission electron microscope (JEOL, Freising, Germany) equipped with a Gatan Orius 1200A camera (Gatan, Munich, Germany).

After prescreening the 250 nm semi-thin sections from HPF samples for ribbon synapses, electron tomography was performed as described previously (Strenzke et al., 2016). First, 10 nm gold beads (British Bio Cell) were applied to both sides of the post-stained grids functioning as fiducial markers. With the Serial-EM software (Mastronarde, 2005), single axis tilt series were acquired mainly from -60° to +60° with 1° increments at 12,000-x magnification and a pixel size of 1.19 nm using a JEM2100 (JEOL) transmission electron microscope at 200 kV. For final tomogram generation, the IMOD software package etomo was used and tomographic reconstructions were generated using 3dmod (Kremer et al., 1996).

### Patch-clamp

Apical turns of the organs of Corti from 2-4 weeks old rats were isolated in ice-cold HEPES Hank’s solution containing (in mM): 5.26 KCl, 141.7 NaCl, 0.5 MgSO_4_.7H_2_O, 10 HEPES, 1 MgCl_2_, 11.1 D-glucose and 3.42 L-glutamine, pH adjusted to around 7.2, osmolality ∼300 mOsm/kg. The recording chamber was perfused with modified Ringer’s solution containing (in mM): 2.8 KCl, 111 NaCl, 35 TEA-Cl, 10 HEPES, 1 CsCl, 1 MgCl_2_, 11.1 D-glucose, 5mM CaCl_2_, pH adjusted to around 7.2 with NaOH and osmolality ∼300 mOsm/kg. Cleaning of the tissue was performed to make the inner hair cells accessible for patch clamp by removing the tectorial membrane and surrounding cells. The clean exposed basolateral surface of IHCs was patch-clamped using EPC-10 amplifier (HEKA Electronics, Germany) controlled by *Patchmaster* software at RT, as also described previously (Moser and Beutner, 2000). Pipettes solutions contained (in mM): 137 Cs-gluconate, 10 TEA-Cl, 10 4-aminopyridine, 10 HEPES, 1 MgCl_2_, and 300µg/ml of amphotericin B, pH adjusted to 7.2 using HCl and CsOH and osmolality ∼290 mOsm/kg.

Cells were kept at a holding potential of -87 mV. All voltages were corrected for liquid junction potential offline (17 mV). Currents were leak corrected using a p/10 protocol. Recordings were discarded when the leak current exceeded -50 pA, Rs exceeded 30 MΩ or Ca^2+^ current rundown exceeded 25%. Current-voltage relationships (IVs) were recorded once the access resistance dropped below 30 MΩ, by applying increasing 10 ms long step-depolarization pulses of voltage ranging from -87 mV to 65 mV, in steps of 5 mV. Exocytosis measurements were performed by measuring increments in membrane capacitance using the Lindau-Neher technique (Lindau and Neher, 1988). Membrane capacitance changes (ΔC_m_) were recorded by stimulating the cells at the potential for maximal Ca^2+^ influx (-17 mV) for variable durations. An interval of 10-90 s was given before the successive stimuli were used. Each protocol was sequentially applied two-three times and only IHCs with reproducible measurements were included. For analysis, traces were averaged 400 ms before and after the depolarisation (skipping the first 60 ms; or first 5, 10, 25 ms in the case of dual pulse experiments with inter-pulse intervals of 25, 50, 100 ms respectively). The traces were subjected to 5 or 10 pass binomial smoothing using Igor Pro 6 (WaveMetrics Inc.) for display.

### Data analysis and statistics

#### Quantitative analysis of FIB-SEM and tomogram datasets

Data were segmented semi-automatically using 3dmod of the IMOD software (Kremer et al., 1996). IHCs, nuclei, afferent nerve fibers, ribbon synapses, mitochondria, pre-and postsynaptic densities were assigned as “closed” objects using the *sculpt* drawing tool. For SVs, first the total amount of vesicles ≤ 80 nm from the ribbon surface were quantified in number and size using the spherical “scattered” object at the maximum projection of the vesicle for spherical SVs. Non-spherical SVs were segmented manually as “closed” objects. The number of SVs was presented as total number per ribbon as well as normalized to the size of the ribbon area. Further, two distinct morphological vesicle pools were additionally quantified in size and number (as previously characterized in (Strenzke et al., 2016)): (i) membrane-proximal synaptic vesicles (MP-SVs, ≤ 50 nm distance between SV membrane and AZ membrane and ≤ 100 nm from the presynaptic density); and (ii) ribbon-associated synaptic vesicles (RA-SVs, first layer of vesicles around the ribbon with a maximum distance of 80 nm from the ribbon surface to the vesicle membrane and not falling into the MP-SV pool). Using the *imodinfo* function of 3dmod, information about the ribbon, mitochondria, pre-and postsynaptic density sizes was given as well as the radii for all vesicle pools were determined with this function in order to calculate the average diameter per tomogram. Distance measurements were performed with the measurement drawing tool along the x, y and z-axis. Movies were generated with 3dmod, Fiji (Schindelin et al., 2012) and iMovie (Apple Inc., version 10.3.1).

#### Quantitative analysis of confocal and STED images

The size of the synaptic ribbon, PSD and Ca^2+^ channel clusters (lines and spots) were estimated by fitting a 2D-Gaussian function to individual structures in 2D-STED or confocal images. This yielded values of full width of half maximum (FWHM) along the long and short axes. The areas of the structures have been reported as areas of ellipsoids, calculated as: Area = π X (Long Axis/2) X (Short Axis/2). For estimation of centers of mass of the immunofluorescent spots, the following formula was used:

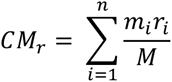

Where *CM_r_* represents the xy coordinates of the center of mass, *m_i_* is the intensity of individual pixel, *r_i_* is the xy coordinate of the pixel and *M* is the sum of all intensities.

All images were analyzed and z-projected using NIH ImageJ software, and further analysis was performed using Igor Pro 6 (WaveMetrics Inc.). For analysis of nearest neighbor distance of synaptic ribbons from immunolabelled confocal sections, we used *Imaris 9.6* (Oxford Instruments). Immunofluorescent ribbon spots were detected per IHC using the inbuilt spot-detection algorithm in a region of interest marked within a z-stack. Spot parameters provided for ellipsoid detection were 0.44 µm and 1.13µm for estimated XY and Z diameters respectively. Thresholding was performed based on quality of immunofluorescence to eliminate artefactual spots. A manual check was performed to include undetected spots and to exclude falsely detected spots and spots not localizing within the cell of interest. The nearest neighbor distance was computed as the minimum distance between the centers of homogenous mass of the detected spots.

#### Statistical analysis

Data are mainly presented as box plots with the mean values highlighted as a cross and with individual data points overlaid and were analysed using Excel, Igor Pro 6 and 7 (WaveMetrics Inc.) and R. In order to characterize the subgroups of *Pclo^gt/gt^* ribbon synapses, acquired with electron tomography, in addition to our manual grouping, we performed an unsupervised K-means clustering with R by considering the following variables: ribbon, presynaptic density as well as SV pool counts and sizes. An optimal number of k=2 clusters was selected based on the decrease of the total within-cluster sum of squares observed.

Using Igor Pro, normality of data was assessed with the Jarque-Bera test or the Wald-Wolfowitz runs test and equality of variances in normally distributed data was assessed with the F-test. Differences between two groups were evaluated for significant differences using the two-tailed unpaired Student’s t-test, or, when not normally distributed and/or variance was unequal, the unpaired two-tailed Mann– Whitney Wilcoxon test. For multiple comparisons, the one-way ANOVA with post-hoc Tukey’s test or the Kruskal-Wallis (KW) test with multiple comparison correction (NPMC: Non-parametric multiple comparison test) was utilized. For Fig 1B, C, we performed two-way repeated measures ANOVA with post-hoc Holm-Šidák correction for multiple comparisons using *GraphPad Prism* 9 (GraphPad Software). Non-significant differences between samples are indicated as *n.s.*, significant differences are indicated as *\*P* < 0.05, *\*\*P* < 0.01, *\*\*\*P* < 0.001, *\*\*\*\*P* < 0.0001.

## Acknowledgements

We thank S. Gerke, C. Senger-Freitag, A.J. Goldak, S. Langer, S. Thom for expert technical assistance. Further, we would like to thank N. Karagulyan for help with analysis scripts.

## Authors contributions

S.M., R.K., T.M. and C.W. designed the study. S.M. performed electron microscopic work (conventional embeddings, HPF/FS, enhanced en bloc stainings, TEM of random sections, electron tomography, FIB-SEM data acquisition and post-processing) with contribution from C.W., data analysis and supervised I.F. together with C.W. for pre-embedding immunogold labeling data acquisition. R.K. performed immunohistochemistry, confocal and STED microscopy, systems physiology, physiological cell data acquisition as well as data analysis with contribution from J.N.. A.M.S. performed and W.M. supervised FIB-SEM data acquisition and post-processing. J.N. supervised electrophysiology. F.A., F.K.H and C.C.G provided unpublished material and contributed to pilot experiment. N.S. supervised systems physiology data acquisition and analysis. M.G. contributed to statistical analysis. S.M., R.K., T.M. and C.W. prepared the manuscript with contributions from all other authors.

## References

1. Ackermann, F., Schink, K. O., Bruns, C., Izsvák, Z., Hamra, F. K., Rosenmund, C. and Garner, C. C. (2019). Critical role for Piccolo in synaptic vesicle retrieval. eLife 8, e46629.

2. Altrock, W. D., tom Dieck, S., Sokolov, M., Meyer, A. C., Sigler, A., Brakebusch, C., Fässler, R., Richter, K., Boeckers, T. M., Potschka, H., et al. (2003). Functional Inactivation of a Fraction of Excitatory Synapses in Mice Deficient for the Active Zone Protein Bassoon. Neuron 37, 787–800.

3. Becker, L., Schnee, M. E., Niwa, M., Sun, W., Maxeiner, S., Talaei, S., Kachar, B., Rutherford, M. A. and Ricci, A. J. (2018). The presynaptic ribbon maintains vesicle populations at the hair cell afferent fiber synapse. eLife 7, e30241.

4. Buran, B. N., Strenzke, N., Neef, A., Gundelfinger, E. D., Moser, T. and Liberman, M. C. (2010). Onset coding is degraded in auditory nerve fibers from mutant mice lacking synaptic ribbons. J. Neurosci. Off. J. Soc. Neurosci. 30, 7587–7597.

5. Butola, T., Wichmann, C. and Moser, T. (2017). Piccolo Promotes Vesicle Replenishment at a Fast Central Auditory Synapse. Front. Synaptic Neurosci. 9, 14.

6. Cases-Langhoff, C., Voss, B., Garner, A. M., Appeltauer, U., Takei, K., Kindler, S., Veh, R. W., De Camilli, P., Gundelfinger, E. D. and Garner, C. C. (1996). Piccolo, a novel 420 kDa protein associated with the presynaptic cytomatrix. Eur. J. Cell Biol. 69, 214–223.

7. Chakrabarti, R. and Wichmann, C. (2019). Nanomachinery Organizing Release at Neuronal and Ribbon Synapses. Int. J. Mol. Sci. 20, 2147.

8. Chakrabarti, R., Michanski, S. and Wichmann, C. (2018). Vesicle sub-pool organization at inner hair cell ribbon synapses. EMBO Rep. e44937.

9. Chapochnikov, N. M., Takago, H., Huang, C.-H., Pangršič, T., Khimich, D., Neef, J., Auge, E., Göttfert, F., Hell, S. W., Wichmann, C., et al. (2014). Uniquantal Release through a Dynamic Fusion Pore Is a Candidate Mechanism of Hair Cell Exocytosis. Neuron 17, 1389–1403.

10. Deerinck, T. J., Shone, T. M., Bushong, E. A., Ramachandra, R., Peltier, S. T. and Ellisman, M. H. (2018). High-performance serial block-face SEM of nonconductive biological samples enabled by focal gas injection-based charge compensation. J. Microsc. 270, 142–149.

11. Dick, O., Hack, I., Altrock, W. D., Garner, C. C., Gundelfinger, E. D. and Brandstätter, J. H. (2001). Localization of the presynaptic cytomatrix protein Piccolo at ribbon and conventional synapses in the rat retina: Comparison with Bassoon. J. Comp. Neurol. 439, 224–234.

12. Dick, O., tom Dieck, S., Altrock, W. D., Ammermüller, J., Weiler, R., Garner, C. C., Gundelfinger, E. D. and Brandstätter, J. H. (2003). The Presynaptic Active Zone Protein Bassoon Is Essential for Photoreceptor Ribbon Synapse Formation in the Retina. Neuron 37, 775–786.

13. Fenster, S. D., Kessels, M. M., Qualmann, B., Chung, W. J., Nash, J., Gundelfinger, E. D. and Garner, C. C. (2003). Interactions between Piccolo and the Actin/Dynamin-binding Protein Abp1 Link Vesicle Endocytosis to Presynaptic Active Zones. J. Biol. Chem. 278, 20268–20277.

14. Frank, T., Khimich, D., Neef, A. and Moser, T. (2009). Mechanisms contributing to synaptic Ca2+ signals and their heterogeneity in hair cells. Proc. Natl. Acad. Sci. 106, 4483–4488.

15. Frank, T., Rutherford, M. A., Strenzke, N., Neef, A., Pangršič, T., Khimich, D., Fejtova, A., Gundelfinger, E. D., Liberman, M. C., Harke, B., et al. (2010). Bassoon and the synaptic ribbon organize Ca^2+^ channels and vesicles to add release sites and promote refilling. Neuron 68, 724–738.

16. Goutman, J. D. and Glowatzki, E. (2007). Time course and calcium dependence of transmitter release at a single ribbon synapse. Proc. Natl. Acad. Sci. 104, 16341–16346.

17. Grabner, C. P. and Moser, T. (2021). The mammalian rod synaptic ribbon is essential for Cav channel facilitation and ultrafast synaptic vesicle fusion. eLife 10, e63844.

18. Grabner, C. P., Gandini, M. A., Rehak, R., Le, Y., Zamponi, G. W. and Schmitz, F. (2015). RIM1/2-Mediated Facilitation of Cav1.4 Channel Opening Is Required for Ca^2+^-Stimulated Release in Mouse Rod Photoreceptors. J. Neurosci. 35, 13133–13147.

19. Gundelfinger, E. D., Reissner, C. and Garner, C. C. (2015). Role of Bassoon and Piccolo in Assembly and Molecular Organization of the Active Zone. Front. Synaptic Neurosci. 7, 19.

20. Hua, Y., Ding, X., Wang, H., Wang, F., Lu, Y., Neef, J., Gao, Y., Moser, T. and Wu, H. (2021). Electron Microscopic Reconstruction of Neural Circuitry in the Cochlea. Cell Rep. 34,.

21. Hull, C., Studholme, K., Yazulla, S. and von Gersdorff, H. (2006). Diurnal changes in exocytosis and the number of synaptic ribbons at active zones of an ON-type bipolar cell terminal. J. Neurophysiol. 96, 2025–2033.

22. Jean, P., Morena, D. L. de la Michanski, S., Tobón, L. M. J., Chakrabarti, R., Picher, M. M., Neef, J., Jung, S., Gültas, M., Maxeiner, S., et al. (2018). The synaptic ribbon is critical for sound encoding at high rates and with temporal precision. eLife 7, e29275.

23. Jean, P., Anttonen, T., Michanski, S., Diego, A. M. G. de Steyer, A. M., Neef, A., Oestreicher, D., Kroll, J., Nardis, C., Pangršič, T., et al. (2020). Macromolecular and electrical coupling between inner hair cells in the rodent cochlea. Nat. Commun. 11, 1–14.

24. Jing, Z., Rutherford, M. A., Takago, H., Frank, T., Fejtova, A., Khimich, D., Moser, T. and Strenzke, N. (2013). Disruption of the presynaptic cytomatrix protein bassoon degrades ribbon anchorage, multiquantal release, and sound encoding at the hair cell afferent synapse. J. Neurosci. 33, 4456–4467.

25. Jung, S., Oshima-Takago, T., Chakrabarti, R., Wong, A. B., Jing, Z., Yamanbaeva, G., Picher, M. M., Wojcik, S. M., Göttfert, F., Predoehl, F., et al. (2015a). Rab3-interacting molecules 2α and 2β promote the abundance of voltage-gated CaV1.3 Ca^2+^ channels at hair cell active zones. Proc. Natl. Acad. Sci. 112, E3141–E3149.

26. Jung, S., Maritzen, T., Wichmann, C., Jing, Z., Neef, A., Revelo, N. H., Al-Moyed, H., Meese, S., Wojcik, S. M., Panou, I., et al. (2015b). Disruption of adaptor protein 2μ (AP-2μ) in cochlear hair cells impairs vesicle reloading of synaptic release sites and hearing. EMBO J. 34, 2686–2702.

27. Kantardzhieva, A. V., Liberman, M. C. and Sewell, W. F. (2013). Quantitative analysis of ribbons, vesicles, and cisterns at the cat inner hair cell synapse: correlations with spontaneous rate. J. Comp. Neurol. 521, 3260–3271.

28. Khimich, D., Nouvian, R., Pujol, R., tom Dieck, S., Egner, A., Gundelfinger, E. D. and Moser, T. (2005). Hair cell synaptic ribbons are essential for synchronous auditory signalling. Nature 434, 889– 894.

29. Kiang, N. Y., Pfeiffer, R. R., Warr, W. B. and Backus, A. S. (1965). Stimulus coding in the cochlear nucleus. Trans. Am. Otol. Soc. 53, 35–58.

30. Kremer, J. R., Mastronarde, D. N. and McIntosh, J. R. (1996). Computer visualization of three-dimensional image data using IMOD. J. Struct. Biol. 116, 71–76.

31. Krinner, S., Butola, T., Jung, S., Wichmann, C. and Moser, T. (2017). RIM-Binding Protein 2 Promotes a Large Number of CaV1.3 Ca^2+^-Channels and Contributes to Fast Synaptic Vesicle Replenishment at Hair Cell Active Zones. Front. Cell. Neurosci. 11, 334.

32. Kroll, J., Jaime Tobón, L. M., Vogl, C., Neef, J., Kondratiuk, I., König, M., Strenzke, N., Wichmann, C., Milosevic, I. and Moser, T. (2019). Endophilin-A regulates presynaptic Ca^2+^ influx and synaptic vesicle recycling in auditory hair cells. EMBO J. 38,.

33. Leal-Ortiz, S., Waites, C. L., Terry-Lorenzo, R., Zamorano, P., Gundelfinger, E. D. and Garner, C. C. (2008). Piccolo modulation of Synapsin1a dynamics regulates synaptic vesicle exocytosis. J. Cell Biol. 181, 831–846.

34. Li, C., Li, X., Bi, Z., Sugino, K., Wang, G., Zhu, T. and Liu, Z. (2020). Comprehensive transcriptome analysis of cochlear spiral ganglion neurons at multiple ages. eLife 9,.

35. Li, P., Lin, Z., An, Y., Lin, J., Zhang, A., Wang, S., Tu, H., Ran, J., Wang, J., Liang, Y., et al. (2021). Piccolo is essential for the maintenance of mouse retina but not cochlear hair cell function. Aging 13, 11678–11695.

36. Liberman, M. C. (1978). Auditory-nerve response from cats raised in a low-noise chamber. J. Acoust. Soc. Am. 63, 442–455.

37. Liberman, M. C. (1980). Morphological differences among radial afferent fibers in the cat cochlea: an electron-microscopic study of serial sections. Hear. Res. 3, 45–63.

38. Liberman, M. C., Dodds, L. W. and Pierce, S. (1990). Afferent and efferent innervation of the cat cochlea: quantitative analysis with light and electron microscopy. J. Comp. Neurol. 301, 443– 460.

39. Liberman, L. D., Wang, H. and Liberman, M. C. (2011). Opposing Gradients of Ribbon Size and AMPA Receptor Expression Underlie Sensitivity Differences among Cochlear-Nerve/Hair-Cell Synapses. J. Neurosci. 31, 801–808.

40. Limbach, C., Laue, M. M., Wang, X., Hu, B., Thiede, N., Hultqvist, G. and Kilimann, M. W. (2011). Molecular in situ topology of Aczonin/Piccolo and associated proteins at the mammalian neurotransmitter release site. Proc. Natl. Acad. Sci. 108, E392–E401.

41. Lindau, M. and Neher, E. (1988). Patch-clamp techniques for time-resolved capacitance measurements in single cells. Pflüg. Arch. Eur. J. Physiol. 411, 137–146.

42. Mastronarde, D. N. (2005). Automated electron microscope tomography using robust prediction of specimen movements. J. Struct. Biol. 152, 36–51.

43. Matthews, G. and Fuchs, P. (2010). The diverse roles of ribbon synapses in sensory neurotransmission. Nat. Rev. Neurosci. 11, 812–822.

44. Maxeiner, S., Luo, F., Tan, A., Schmitz, F. and Südhof, T. C. (2016). How to make a synaptic ribbon: RIBEYE deletion abolishes ribbons in retinal synapses and disrupts neurotransmitter release. EMBO J. 35, 1098–1114.

45. Medrano, G. A., Singh, M., Plautz, E. J., Good, L. B., Chapman, K. M., Chaudhary, J., Jaichander, P., Powell, H. M., Pudasaini, A., Shelton, J. M., et al. (2020). Mutant screen for reproduction unveils depression-associated Piccolo’s control over reproductive behavior. 405985.

46. Meyer, A. C., Frank, T., Khimich, D., Hoch, G., Riedel, D., Chapochnikov, N. M., Yarin, Y. M., Harke, B., Hell, S. W., Egner, A., et al. (2009). Tuning of synapse number, structure and function in the cochlea. Nat. Neurosci. 12, 444–453.

47. Michanski, S., Smaluch, K., Steyer, A. M., Chakrabarti, R., Setz, C., Oestreicher, D., Fischer, C., Möbius, W., Moser, T., Vogl, C., et al. (2019). Mapping developmental maturation of inner hair cell ribbon synapses in the apical mouse cochlea. Proc. Natl. Acad. Sci. 116, 6415–6424.

48. Moser, T. and Beutner, D. (2000). Kinetics of exocytosis and endocytosis at the cochlear inner hair cell afferent synapse of the mouse. Proc. Natl. Acad. Sci. 97, 883–888.

49. Moser, T., Grabner, C. P. and Schmitz, F. (2019). Sensory processing at ribbon synapses in the retina and the cochlea. Physiol. Rev. 100, 103–144.

50. Mukherjee, K., Yang, X., Gerber, S. H., Kwon, H.-B., Ho, A., Castillo, P. E., Liu, X. and Südhof, T. C. (2010). Piccolo and bassoon maintain synaptic vesicle clustering without directly participating in vesicle exocytosis. Proc. Natl. Acad. Sci. U. S. A. 107, 6504–6509.

51. Müller, T. M., Gierke, K., Joachimsthaler, A., Sticht, H., Izsvák, Z., Hamra, F. K., Fejtová, A., Ackermann, F., Garner, C. C., Kremers, J., et al. (2019). A multiple Piccolino-RIBEYE interaction supports plate-shaped synaptic ribbons in retinal neurons. J. Neurosci. 39, 2038–18.

52. Neef, J., Urban, N. T., Ohn, T.-L., Frank, T., Jean, P., Hell, S. W., Willig, K. I. and Moser, T. (2018). Quantitative optical nanophysiology of Ca^2+^ signaling at inner hair cell active zones. Nat. Commun. 9, 290.

53. Ohn, T.-L., Rutherford, M. A., Jing, Z., Jung, S., Duque-Afonso, C. J., Hoch, G., Picher, M. M., Scharinger, A., Strenzke, N. and Moser, T. (2016). Hair cells use active zones with different voltage dependence of Ca^2+^ influx to decompose sounds into complementary neural codes. Proc. Natl. Acad. Sci. 113, E4716–E4725.

54. Özçete, Ö. D. and Moser, T. (2021). A sensory cell diversifies its output by varying Ca^2+^ influx-release coupling among active zones. EMBO J. 40, e106010.

55. Pangrsic, T., Singer, J. H. and Koschak, A. (2018). Voltage-Gated Calcium Channels: Key Players in Sensory Coding in the Retina and the Inner Ear. Physiol. Rev. 98, 2063–2096.

56. Parthier, D., Kuner, T. and Körber, C. (2018). The presynaptic scaffolding protein Piccolo organizes the readily releasable pool at the calyx of Held. J. Physiol. 596, 1485–1499.

57. Payne, S. A., Joens, M. S., Chung, H., Skigen, N., Frank, A., Gattani, S., Vaughn, K., Schwed, A., Nester, M., Bhattacharyya, A., et al. (2021). Maturation of Heterogeneity in Afferent Synapse Ultrastructure in the Mouse Cochlea. Front. Synaptic Neurosci. 0,.

58. Petitpré, C., Wu, H., Sharma, A., Tokarska, A., Fontanet, P., Wang, Y., Helmbacher, F., Yackle, K., Silberberg, G., Hadjab, S., et al. (2018). Neuronal heterogeneity and stereotyped connectivity in the auditory afferent system. Nat. Commun. 9, 3691.

59. Regus-Leidig, H., Ott, C., Löhner, M., Atorf, J., Fuchs, M., Sedmak, T., Kremers, J., Fejtová, A., Gundelfinger, E. D. and Brandstätter, J. H. (2013). Identification and Immunocytochemical Characterization of Piccolino, a Novel Piccolo Splice Variant Selectively Expressed at Sensory Ribbon Synapses of the Eye and Ear. PLoS ONE 8, e70373.

60. Regus-Leidig, H., Fuchs, M., Löhner, M., Leist, S. R., Leal-Ortiz, S., Chiodo, V. A., Hauswirth, W. W., Garner, C. C. and Brandstätter, J. H. (2014). In vivo knockdown of Piccolino disrupts presynaptic ribbon morphology in mouse photoreceptor synapses. Front. Cell. Neurosci. 8, 259.

61. Reynolds, E. S. (1963). THE USE OF LEAD CITRATE AT HIGH pH AS AN ELECTRON-OPAQUE STAIN IN ELECTRON MICROSCOPY. J. Cell Biol. 17, 208–212.

62. Ruel, J., Nouvian, R., D’Aldin, C. G., Pujol, R., Eybalin, M. and Puel, J.-L. (2001). Dopamine inhibition of auditory nerve activity in the adult mammalian cochlea. Eur. J. Neurosci. 14, 977–986.

63. Rutherford, M. A. and Moser, T. (2016). The Ribbon Synapse Between Type I Spiral Ganglion Neurons and Inner Hair Cells. In The Primary Auditory Neurons of the Mammalian Cochlea (ed. Dabdoub, A.), Fritzsch, B.), Popper, A. N.), and Fay, R. R.), pp. 117–156. New York, NY: Springer New York.

64. Sachs, M. B. and Abbas, P. J. (1974). Rate versus level functions for auditory-nerve fibers in cats: tone-burst stimuli. J. Acoust. Soc. Am. 56, 1835–1847.

65. Schindelin, J., Arganda-Carreras, I., Frise, E., Kaynig, V., Longair, M., Pietzsch, T., Preibisch, S., Rueden, C., Saalfeld, S., Schmid, B., et al. (2012). Fiji: an open-source platform for biological-image analysis. Nat. Methods 9, 676–682.

66. Schmitz, F., Königstorfer, A. and Südhof, T. C. (2000). RIBEYE, a component of synaptic ribbons: a protein’s journey through evolution provides insight into synaptic ribbon function. Neuron 28, 857–872.

67. Sherrill, H. E., Jean, P., Driver, E. C., Sanders, T. R., Fitzgerald, T. S., Moser, T. and Kelley, M. W. (2019). Pou4f1 Defines a Subgroup of Type I Spiral Ganglion Neurons and Is Necessary for Normal Inner Hair Cell Presynaptic Ca^2+^ Signaling. J. Neurosci. 39, 5284–5298.

68. Shrestha, B. R., Chia, C., Wu, L., Kujawa, S. G., Liberman, M. C. and Goodrich, L. V. (2018). Sensory Neuron Diversity in the Inner Ear Is Shaped by Activity. Cell 174, 1229–1246.e17.

69. Snellman, J., Mehta, B., Babai, N., Bartoletti, T. M., Akmentin, W., Francis, A., Matthews, G., Thoreson, W. and Zenisek, D. (2011). Acute destruction of the synaptic ribbon reveals a role for the ribbon in vesicle priming. Nat. Neurosci. 14, 1135–1141.

70. Sobkowicz, H. M., Rose, J. E., Scott, G. E. and Slapnick, S. M. (1982). Ribbon synapses in the developing intact and cultured organ of Corti in the mouse. J. Neurosci. 2, 942–957.

71. Sobkowicz, H. M., Rose, J. E., Scott, G. L. and Levenick, C. V. (1986). Distribution of synaptic ribbons in the developing organ of Corti. J. Neurocytol. 15, 693–714.

72. Stamataki, S., Francis, H. W., Lehar, M., May, B. J. and Ryugo, D. K. (2006). Synaptic alterations at inner hair cells precede spiral ganglion cell loss in aging C57BL/6J mice. Hear. Res. 221, 104–118.

73. Strenzke, N., Chakrabarti, R., Al-Moyed, H., Müller, A., Hoch, G., Pangrsic, T., Yamanbaeva, G., Lenz, C., Pan, K.-T., Auge, E., et al. (2016). Hair cell synaptic dysfunction, auditory fatigue and thermal sensitivity in otoferlin Ile515Thr mutants. EMBO J. 35, e201694564.

74. Sun, S., Babola, T., Pregernig, G., So, K. S., Nguyen, M., Su, S.-S. M., Palermo, A. T., Bergles, D. E., Burns, J. C. and Müller, U. (2018). Hair Cell Mechanotransduction Regulates Spontaneous Activity and Spiral Ganglion Subtype Specification in the Auditory System. Cell 174, 1247–1263.e15.

75. Taberner, A. M. and Liberman, M. C. (2005). Response Properties of Single Auditory Nerve Fibers in the Mouse. J. Neurophysiol. 93, 557–569.

76. tom Dieck, S., Sanmartí-Vila, L., Langnaese, K., Richter, K., Kindler, S., Soyke, A., Wex, H., Smalla, K. H., Kämpf, U., Fränzer, J. T., et al. (1998). Bassoon, a novel zinc-finger CAG/glutamine-repeat protein selectively localized at the active zone of presynaptic nerve terminals. J. Cell Biol. 142, 499–509.

77. Waites, C. L., Leal-Ortiz, S. A., Okerlund, N., Dalke, H., Fejtova, A., Altrock, W. D., Gundelfinger, E. D. and Garner, C. C. (2013). Bassoon and Piccolo maintain synapse integrity by regulating protein ubiquitination and degradation. EMBO J. 32, 954–969.

78. Wichmann, C. and Moser, T. (2015). Relating structure and function of inner hair cell ribbon synapses. Cell Tissue Res. 361, 95–114.

79. Winter, I. M., Robertson, D. and Yates, G. K. (1990). Diversity of characteristic frequency rate-intensity functions in guinea pig auditory nerve fibres. Hear. Res. 45, 191–202.

80. Wong, A. B., Rutherford, M. A., Gabrielaitis, M., Pangršič, T., Göttfert, F., Frank, T., Michanski, S., Hell, S., Wolf, F., Wichmann, C., et al. (2014). Developmental refinement of hair cell synapses tightens the coupling of Ca^2+^ influx to exocytosis. EMBO J. 33, 247–264.

81. Yin, Y., Liberman, L. D., Maison, S. F. and Liberman, M. C. (2014). Olivocochlear innervation maintains the normal modiolar-pillar and habenular-cuticular gradients in cochlear synaptic morphology. J. Assoc. Res. Otolaryngol. JARO 15, 571–583.

